# ERK pathway activation inhibits ciliogenesis and causes defects in motor behavior, ciliary gating, and cytoskeletal rearrangement

**DOI:** 10.1101/2022.05.18.492507

**Authors:** Larissa L Dougherty, Soumita Dutta, Prachee Avasthi

**Author notes:** Address correspondence to at Geisel School of Medicine at Dartmouth College. Prachee Avasthi is a paid consultant for Arcadia Science.

## Abstract

Mitogen-activated protein kinase (MAPK) pathways are well known regulators of the cell cycle but they have also been found to control ciliary length in a wide variety of organisms and cell types from *Caenorhabditis elegans* neurons to mammalian photoreceptors through unknown mechanisms. ERK1/2 is a MAP kinase in human cells that is predominantly phosphorylated by MEK1/2 and dephosphorylated by the phosphatase DUSP6. We have found that the ERK1/2 activator/DUSP6 inhibitor, (E)-2-benzylidine-3-(cyclohexylamino)-2,3-dihydro-1H-inden-1-one (BCI), inhibits ciliary maintenance in *Chlamydomonas* and hTERT-RPE1 cells and assembly in *Chlamydomonas*. These effects involve inhibition of total protein synthesis, microtubule organization, membrane trafficking, and partial kinesin-2 motor dynamics. Our data provide evidence for various avenues for BCI-induced ciliary shortening and impaired ciliogenesis that gives mechanistic insight into how MAP kinases can regulate ciliary length.

## INTRODUCTION

### The cell cycle and ciliogenesis utilize the same structures at different times

Proper ciliary function is important for cellular signaling and development in most mammalian cells (Reiter & Leroux, 2017). Dysregulation of these antenna-like microtubule signaling/motile hubs can result in problems ranging from polydactyly to retinal dystrophy (Reiter & Leroux, 2017). Ciliary assembly and maintenance is regulated by intraflagellar transport (IFT) which controls the movement of materials into and out of the cilium (Ishikawa & Marshall, 2017).

Ciliogenesis occurs when cells the exit cell cycle (Kasahara & Inagaki, 2021). After cell division and during quiescence, the centrosome takes on a new role from serving as spindle poles for chromosome segregation to serving as basal bodies at the base of the cilium (Lattao et al., 2017). One of the two centrioles in the centrosome, the mother centriole, docks to the plasma membrane during G1/G0 to nucleate the cilium and recruit proteins involved in ciliogenesis (Azimzadeh & Marshall, 2010). In *Chlamydomonas*, two cilia begin to assemble from microtubule growth pushing from the basal bodies into the plasma membrane. This creates a protrusion which ultimately becomes a cilium. As the cell reenters the cell cycle, the cilium will disassemble either through shedding or axonemal disassembly before entering cell division (Patel & Tsiokas, 2021).

### The ERK pathway controls the cell cycle

MAPK pathways regulate cellular stress responses, developmental phases of cells, and cell proliferation (Zhang & Liu, 2002). The Ras/Raf/MEK/ERK pathway is one of the central/core signaling pathways that has been well characterized for incorporating extracellular signals into the cell to promote cell proliferation and differentiation (Shaul & Seger, 2007). At its most basic regulation in human cells, it involves the balance of ERK1/2 phosphorylation by the kinase MEK1/2, and the phosphatase DUSP6/MKP3 (Lake et al., 2016), though there is extensive crosstalk, other proteins, and feedback loops also involved in regulation of this phosphorylation cascade.

### Various MAPKs have been found to regulate ciliary structure

Prior work has shown that mutants for MAPKs can modulate cilium length. For example, a type of MAPK in *Chlamydomonas*, LF4 which is a MAPK/MAK/MRK Overlapping Kinase (MOK), leads to longer cilia when mutated (Berman et al., 2003) as well as mutated MPK9 in *Leishmania Mexican*a (Bengs et al., 2005). In photoreceptor cells, male germ cell-associated kinase (MAK) negatively regulates ciliary length to prevent degeneration (Kazatskaya et al., 2017). In *C. elegans*, mutations to MAPK15 directly regulate primary cilium formation and localization of the ciliary proteins BBS7 and other proteins involved in cilia formation and maintenance. More recently, ERK7, another MAPK, has been found to regulate an actin regulating protein, CAP Zip, which is necessary for ciliary length maintenance in conjunction with other signaling pathways (Miyatake et al., 2015). In addition, it is known that ERK1/2 suppression with 1,4-diamino-2,3-dicyano-1,4-bis[2-aminophenylthio]butadiene, or U0126, can elongate cilia as well as decrease apoptosis in kidney cells (S. Wang et al., 2013). Our goal in this work is to understand how the central signaling MAPK pathway can regulate ciliogenesis in addition to its role in regulating the cell cycle (Wortzel & Seger, 2011).

Here we use *Chlamydomonas reinhardtii*, a unicellular green alga which is an extensively used ciliary model organism with well conserved ciliary proteins, structure, and function to humans (O’Toole et al., 2012). BCI-induced ERK1/2 activation alters multiple pathways that are important for ciliogenesis and ciliary maintenance. This pathway has been previously manipulated through the MEK1/2 inhibitor U0126 to prevent MAPK activation as well as lengthen cilia (Avasthi et al., 2012). To understand why this process occurs, we looked at various processes that are involved in ciliary length maintenance and assembly.

## RESULTS

### BCI-induced ERK1/2 phosphorylation disrupts ciliary maintenance and assembly in Chlamydomonas reinhardtii

Given the lengthening effects of ERK inhibition on ciliary length through U0126 (Avasthi et al., 2012), we wanted to test specificity of the pathway in ciliary regulation through activation of the pathway. The ERK activator (E)-2-benzylidene-3-(cyclohexylamino)-2,3-dihydro-1H-inden-1-one (BCI) has previously been found to inhibit the dual specificity phosphatase/MAP kinase phosphatase, DUSP6/MKP3 (Molina et al., 2009; Shojaee et al., 2015), and subsequently activate ERK1/2 (**Fig 1A**). To test if this intended effect occurs in *Chlamydomonas*, we checked MAPK phosphorylation following 15 minute intervals of treatment with BCI for up to 1 hour. Phosphorylation increased within 15 minutes and then decreased by 60 minutes, indicating that BCI activates phosphorylation of MAP Kinases in *Chlamydomonas* **(Fig 1B**).

**Figure 1.**
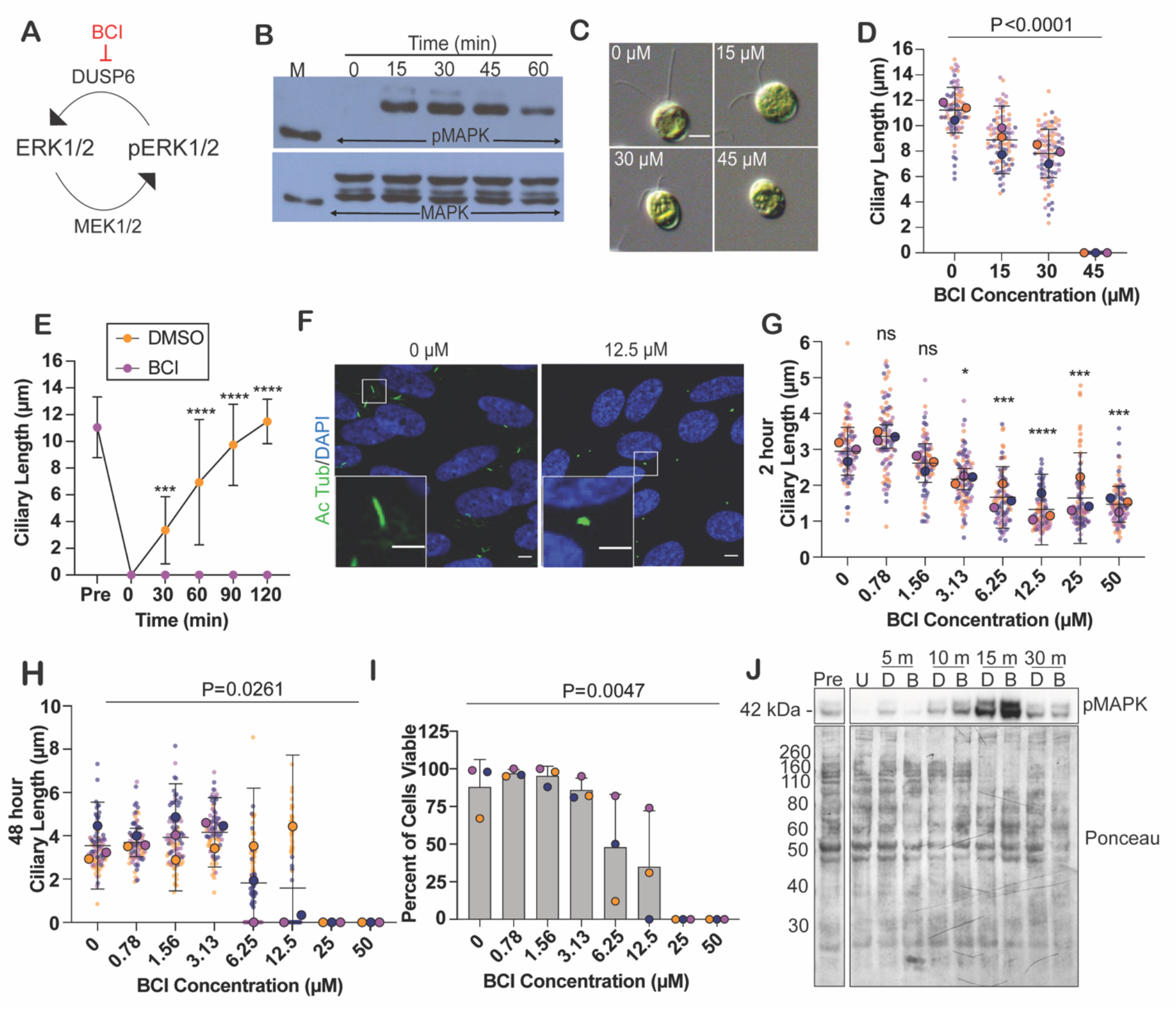
BCI activates the MAPK pathway in *Chlamydomonas reinhardtii* and hTERT-RPE1 and inhibits ciliogenesis. **(A)** The MAPK kinase ERK1/2 is phosphorylated by MEK1/2 and dephosphorylated by DUSP6. **(B)** Western blot showing an increase in pMAPK protein expression following treatment with 30 *µ*M BCI for the indicated times. MAPK protein expression is unchanged. **(C)** Representative images of steady state *Chlamydomonas* cilia length in 0 *µ*M BCI (1% DMSO) to 45 *µ*M BCI. Scale bar is 5 *µ*m. (**D)** Quantification of ciliary lengths in (C) following a 2 hour treatment with the indicated concentrations of BCI. Error bars show the mean with the 95% confidence interval for averages from each trial (n=30, N=30). The P value was calculated with a Brown-Forsythe ANOVA test. This super plot shows averages from each trial with larger circles plotted on top of the individual points. (**E**) Regenerating ciliary lengths following pH shock and regrowth in 30 *µ*M BCI or 0.5% DMSO over 2 hours. Error bars are mean with 95% confidence of the means from each trial (n=30, N=3). The P values for the 120 minute time points were determined using a two-way ANOVA with Bonferroni’s correction (**** P<0.0001, *** P=0.0002). **(F)** Representative images of treated serum starved RPE1 control cells (0.6% DMSO) compared to the shortest cilium lengths measured following a 2 hour BCI treatment (12.5 *µ*M BCI). Cells were stained with acetylated tubulin (green) and DAPI (blue). Acetylated tubulin length was measured for the ciliary length. The scale bar in larger images is 5 *µ*m, and in insets is 3 *µ*m. (**G**) Ciliary lengths of serum starved RPE1 cells treated with the indicated concentrations of BCI for 2 hours. Error bars are mean with 95% confidence interval (n=30, N=3). The P value was calculated with an ordinary one-way ANOVA on the averages of the 3 trials with Bonferonni’s multiple comparisons test. (**H**) RPE1 cells were treated for 48 hours with the indicated concentrations of BCI and then fixed and stained for cilia with acetylated tubulin and DAPI. Ciliary length was measured according to the length of the acetylated tubulin. Error bars are mean with 95% confidence interval (n=30, N=3). P=0.0261 calculated with a Brown-Forsythe ANOVA test. For P values, not significant (ns) is p≥0.05, * is p=0.01 to 0.05, ** is p=0.001 to 0.01, *** is p=0.0001 to 0.001, and **** is p<0.0001. **(I**) Cell viability after 48 hours of serum starvation in combination with various concentrations of BCI. Hoechst stain was used to measure living cells and Sytox Green to measure dead cells. Error bars are mean with standard deviation (n=100, N=3). The P value was calculated with a Browne-Forsythe ANOVA test. **(J)** Western blot showing that 6.25 *µ*M BCI induces ERK1/2 phosphorylation in RPE1 cells. Cells were first incubated with the MEK1/2 inhibitor U0126 to completely inhibit pMAPK signal, and then cells were treated with 6.25 *µ*M for the indicated times. Ponceau staining was used to measure total protein. U is U0126 (50 *µ*M), D is DMSO (0.6%), and B is BCI (6.25 *µ*M).

To determine if ERK1/2 activation through BCI treatment can shorten cilia, we tested a range of BCI concentrations and measured cilia after a 2 hour treatment when length changes are typically apparent. We saw a dose-dependent effect on cilia length where increasing BCI concentrations decreased cilia length up to 45 *µ*M BCI where cilia were completely resorbed (**Fig 1C-D**). This decreased length could be a result of reduced assembly or increased disassembly (Marshall et al., 2005). To test assembly, we severed cilia with low pH and tested ciliary reassembly (Lefebvre, 1995). In the presence of BCI, cilia could not regenerate at all (**Fig 1E)** confirming that there is a strong assembly defect when MAPK is activated through inhibition of its dephosphorylating enzyme despite the presence of excess ciliary protein available for ciliogenesis (Rosenbaum et al., 1969).

To determine if ciliary shortening is due to inhibitor targeting of DUSP6, we used genetic mutants of potential *Chlamydomonas* DUSP6 orthologs. To identify the *Chlamydomonas* DUSP6, we compared the zebrafish DUSP6 protein to the *Chlamydomonas* genome and took the top 3 most similar proteins to investigate which will be referred to here on as the putative DUSP6 orthologs (**SF1A, B**). We first compared their BCI binding pockets. BCI is predicted to interact with zebrafish Trp 262, Asn 335, and Phe 336 as well as the general acid loop backbone according to docking studies (Molina et al., 2009). The predicted zebrafish BCI binding pocket was well conserved to the putative DUSP6 *Chlamydomonas* orthologs (**SF1B**). We acquired these mutants from the *Chlamydomonas* Resource Center, which contain a cassette insertion in the target protein gene to knock out that protein, and tested their ciliary phenotypes (**SF1C-E**). Comparing steady state ciliary lengths showed that the DUSP6 orthologs *mkp2* and *mkp5* both had shorter than wild type ciliary lengths (**SF 1D**). Upon regenerating cilia in these mutants, MKP2 could not regenerate back to wild type length within 2 hours (**SF 1E**). This data supports that there may be other MKPs in *Chlamydomonas* that are the target for BCI.

### BCI-induced ERK1/2 phosphorylation disrupts ciliary maintenance and assembly in *hTERT-RPE1 cells*

We repeated these experiments in hTERT-RPE1 cells to see if this was a *Chlamydomonas*-specific effect. Like *Chlamydomonas*, RPE1 cells showed a dose-dependent effect on ciliary shortening (**Fig 1F-G**). However, unlike *Chlamydomonas*, RPE1 cells were able to assemble cilia in the presence of BCI except at concentrations which were toxic including 25 and 50 *µ*M after 48 hours of treatment and serum starvation (**Fig 1H-I**). In addition, BCI induces ERK1/2 phosphorylation in RPE1 cells within 15 minutes of treatment (**Fig 1J**). Together, this data shows that BCI induces phosphorylation of MAP kinases during short treatments and prevents ciliary assembly.

### BCI disrupts ciliary kinesin-2 dynamics

Ciliary entry of the anterograde motor, kinesin-2, is known to be required for ciliogenesis (Adams et al., 1982). It is possible that cilia shorten and cannot regenerate in the presence of BCI due to blocked kinesin-2 entry into cilia. To test this hypothesis, we first checked if kinesin-2 was present in the cilium and basal bodies of BCI treated *Chlamydomonas* cells using a GFP-tagged kinesin accessory protein on kinesin-2 (KAP-GFP) (**Fig 2A**). After a 2 hour treatment with BCI, cells with cilia maintained KAP-GFP fluorescence at their basal bodies despite increased concentrations of BCI (**Fig 2B**) though their ciliary fluorescence decreased with ciliary length (**Fig 2C**).

**Figure 2.**
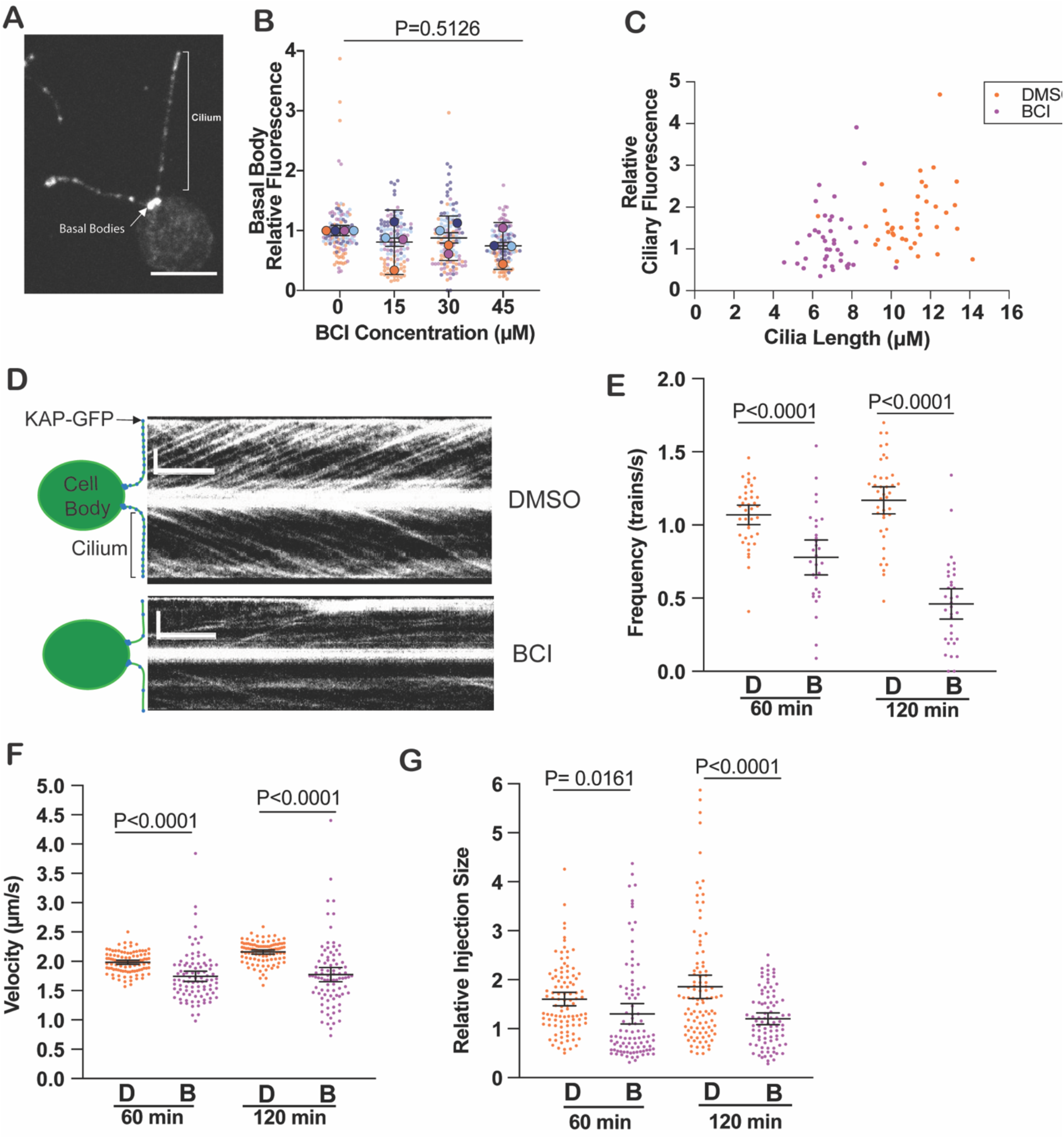
Kinesin-2 dynamics are altered in the cilium of BCI treated cells. (**A**) *Chlamydomonas* expressing KAP-GFP. Indicated are the measured areas of KAP-GFP localization: the 2 basal bodies and cilia. Scale bar is 5 *µ*m. (**B**) Super plot quantification of basal body fluorescence in (A) following a 2 hour treatment with BCI at the indicated concentrations. Error bars are mean with 95% confidence interval. The average from each trial is plotted in larger circles on top of the individual points. P=0.5126 which was determined by an ordinary one-way ANOVA. (**C**) Ciliary length plotted against ciliary fluorescence for cells in DMSO (orange circles) or 30 *µ*M BCI (purple). (**D**) Example kymographs collected from total internal reflection fluorescent (TIRF) microscopy of KAP-GFP movement in cilia in cells treated with DMSO (top) or 30 *µ*M BCI (bottom) for 2 hours. Vertical scale bars are 4 *µ*m. Horizontal scale bars are 2 seconds. (**E-F**) KAP-GFP dynamics quantified from the kymographs (n=20, N=1). The error bars are mean with 95% confidence interval (n=20, N=1). P values were calculated from a two-tailed unpaired t-test for pairwise comparisons. (**E**) Frequency of KAP-GFP trains measured as the total amount of trains counted over the total amount of time the kymograph was collected. For DMSO, n=40 (1 hr and 2 hr). For BCI, n=30 (1 hr) and 34 (2 hr). (**F**) Velocity of KAP-GFP trains measured as the distance traveled in *µ*m over time in seconds. For DMSO, n=100 (1 hr and 2 hr). For BCI, n=93 (1 hr) and 88 (2 hr). (**G**) Relative injection size KAP-GFP trains measured as relative total fluorescent intensity of each train relative to measurements taken before treatment with DMSO or BCI (data not shown). For DMSO, n=100 (1 hr and 2 hr). For BCI, n=93 (1 hr) and 88 (2 hr).

To determine how kinesin-2 dynamics respond to BCI treatment, we used Total Internal Reflection Microscopy (TIRF) to measure kinesin-2 dynamics in real time (**Fig 2D**). Previous studies have shown that kinesin-2 expression increases in shorter regenerating cilia (Brown et al., 1999). From kymographs of KAP-GFP movement in cilia using TIRF, we measured frequency (**Fig 2E**), velocity (**Fig 2F**), and train size (**Fig 2G**) between 60 minutes and 120 minutes. Frequency continually decreased from 60 minutes to 120 minutes in BCI whereas velocity and train size decreased by 60 minutes and maintained that decrease even after 120 minutes. This data suggests that because BCI treatment decreases kinesin-2 entry into cilia, ciliary maintenance cannot be achieved to allow proper ciliogenesis to occur.

### BCI Inhibits kinesin-2 protein synthesis

Decreased KAP-GFP at the basal bodies and cilia could suggest that expression is altered in the whole cell. We checked two concentrations of BCI (**Fig 3A**), 20 *µ*M which induces partial steady state ciliary resorption (**Fig 3B-C**) and 45 *µ*M BCI which induces complete resorption (**Fig 1D, Fig 3D-E**). In partially resorbing cells, cilia have slightly less KAP-GFP though the cell can retain KAP-GFP in the cell body. Similarly, cells with fully resorbed cilia do not show any decrease in expression. We checked this total expression in regenerating cells as well (**Fig 3F-I**). At 0 minutes, kinesin-2 fluorescent intensity at the basal bodies slightly increased and then decreased to pre levels by 2 hours when cilia are almost fully formed; however, in BCI treated cells, this expression did not decrease after two hours (**Fig 3G**) indicating that KAP-GFP can get recruited to the basal bodies for ciliary entry despite not being able to enter. Interestingly, we found that total KAP-GFP protein expression in the cell was not increased in BCI-treated cells after 2 hours (**Fig 3H-I**) which was only partially rescued with the proteasome inhibitor MG132 (carbobenzoxy-Leu-Leu-leucinal) (**SF 2A and B**). This data indicates that BCI inhibits KAP-GFP protein expression in the cell but does not prevent signaling events involved in recruiting KAP-GFP to the cilium.

**Figure 3.**
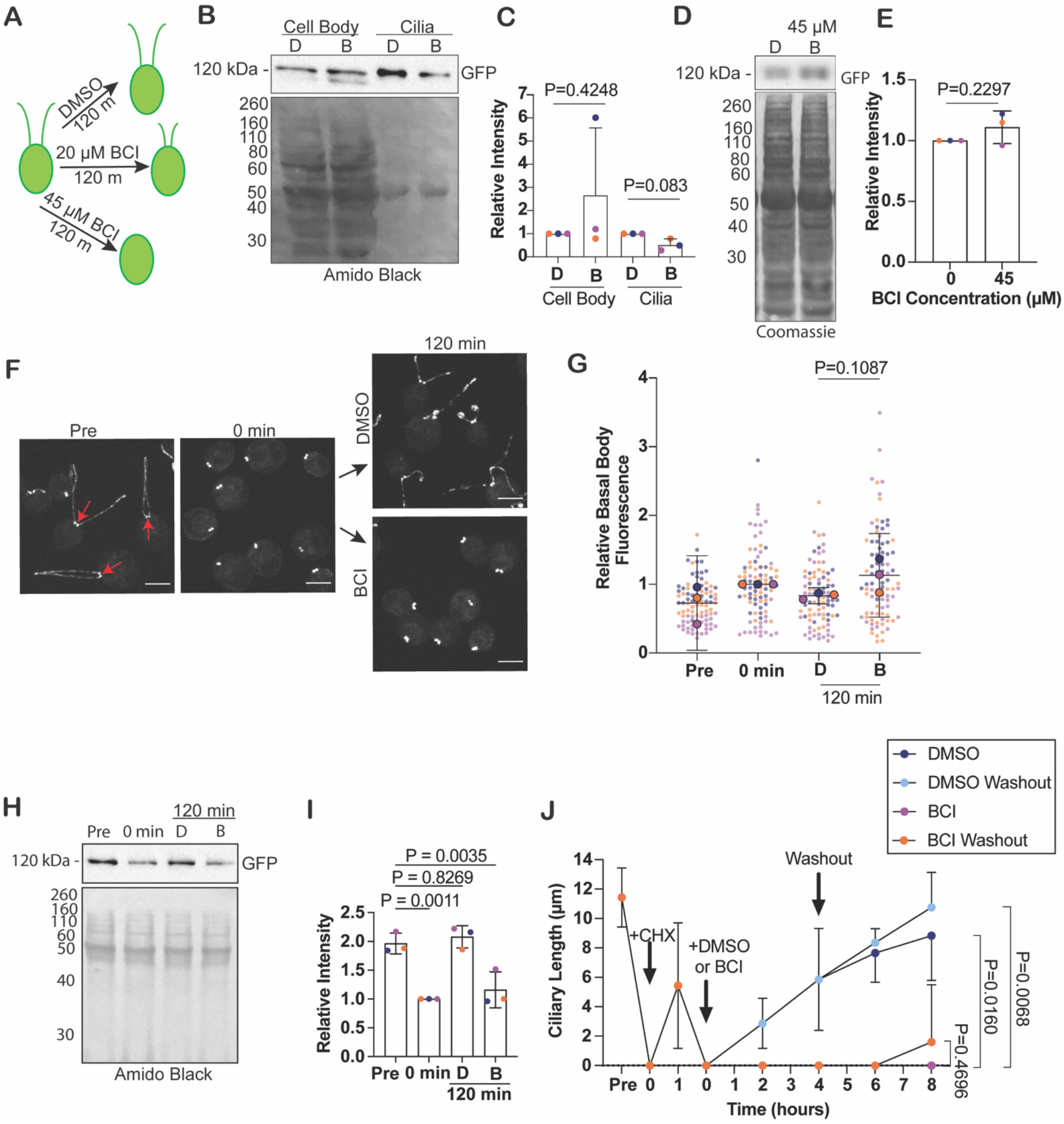
BCI decreases KAP-GFP protein expression without impacting its recruitment/targeting. **(A)** Experimental illustration for (B-E). *Chlamydomonas* cells were treated for 2 hours with either 0.5% DMSO, 30 *µ*M BCI to induce partial ciliary resorption, or 45 *µ*M BCI to induce complete ciliary resorption. **(B)** Western blot for KAP-GFP expression in the presence of DMSO or 30 *µ*M BCI for 2 hours. Cilia were isolated and KAP-GFP expression checked separately from the cell bodies using a GFP antibody. Total protein was determined with amido black. (**C**) Intensity quantification of GFP normalized to total protein. Error bars are standard deviation of the mean. P values were determined by a Welch’s t test (N=3). D is DMSO (0.5%), B is BCI (30 *µ*M). **(D)** Western blot for KAP-GFP expression in the presence of DMSO or 45 *µ*M BCI to induce complete ciliary resorption. GFP antibody was used to probe for KAP-GFP. Total protein was determined with Coomassie Brilliant Blue. (**E**) Intensity quantification (right) measures protein from 3 independent experiments. Error bars are mean with standard deviation. **(F)** Immunofluorescent images of KAP-GFP cells during regeneration in either 0.5% DMSO or 30 *µ*M BCI. Scale bars are 5 *µ*m. Red arrows point to basal bodies which are the object of quantification in (G). **(G)** Quantification of KAP-GFP fluorescence at the basal bodies. Error bars are mean with 95% confidence interval of the averages for 3 independent trials (n=30, N=3). The P value was calculated using an unpaired t-test with Welch’s correction. **(H)** Western blot for KAP-GFP expression in regenerating cells in either DMSO or 30 *µ*M BCI. Total protein was measured with amido black. **(I)** Quantification of F. Error bars are mean with standard deviation for 3 independent experiments. P values were calculated using an ordinary one-way ANOVA with Dunnett’s multiple comparisons test. **(J)** Double deciliation with a 1 hour regrowth in cycloheximide (CHX). After the first pH shock, cycloheximide (CHX) is added for the first 60 minutes. Cells were deciliated once more, regenerated in 0.2% DMSO or 20 *µ*M BCI for 4 hours, and washed out or not and grown for 4 more hours. Error bars are 95% confidence interval on the means of 3 experiments (n=30, N=3). P values were determined with a two-way ANOVA with Tukey’s correction for multiple comparisons.

### BCI Inhibits ciliary protein synthesis

Previous work has shown that there is an existing ciliary protein pool that has enough protein to synthesize *Chlamydomonas* cilia to half length in the absence of protein synthesis with cycloheximide (Lefebvre et al., 1978). We wanted to see what happens to protein synthesis and ciliary incorporation during ciliogenesis in BCI-treated cells. We regenerated cells for 1 hour in cycloheximide to deplete the protein pool and then deciliated them once more and let them regrow in either DMSO or BCI for 4 hours before washing out the drugs to determine if BCI is inhibiting protein synthesis (**Fig 3J**). In BCI over 8 hours, or after a 4 hour washout, cells could not regenerate their cilia which indicates that BCI potently inhibits total ciliary protein synthesis in regenerating cells (**Figure 3J**). Meanwhile, singly regenerating cells in the same concentration of BCI can begin regenerating their cilia slowly 3 hours post deciliation and at the same rate after washing out BCI after 2 hours (**SF 2C**). Together, this data indicates that BCI inhibits ciliary protein synthesis.

### BCI partially disrupts the transition zone

The transition zone is the ciliary gate of the cell which works to regulate protein entry and exit at the cilium (Gonçalves & Pelletier, 2017). Decreased kinesin-2 expression at the basal bodies and within cilia suggest that there could be a defect in the transition zone structure. To test this, we first looked at the structure with electron microscopy. Cross sections of the transition zone in both untreated and BCI-treated cells appeared unaltered, noting the classical “H shape” and wedge connectors (Diener et al., 2015), though it might be apparent that the protein density is more localized to the bottom of the transition zone in BCI-treated cells as opposed to the top (**Fig 4A**). We also checked a transition zone protein, NPHP4, in BCI-treated cells. In steady state and in regenerating cells with BCI, NPHP4 still localized as two puncta at the transition zone which supports the unaltered structure visualized with EM, though NPHP4 immunofluorescence was decreased (**Fig 4B-C**). Together, these data support that BCI does not grossly disrupt the transition zone as a mechanism for altering ciliogenesis, though it may inhibit expression of proteins or protein turnover at the transition zone.

**Figure 4.**
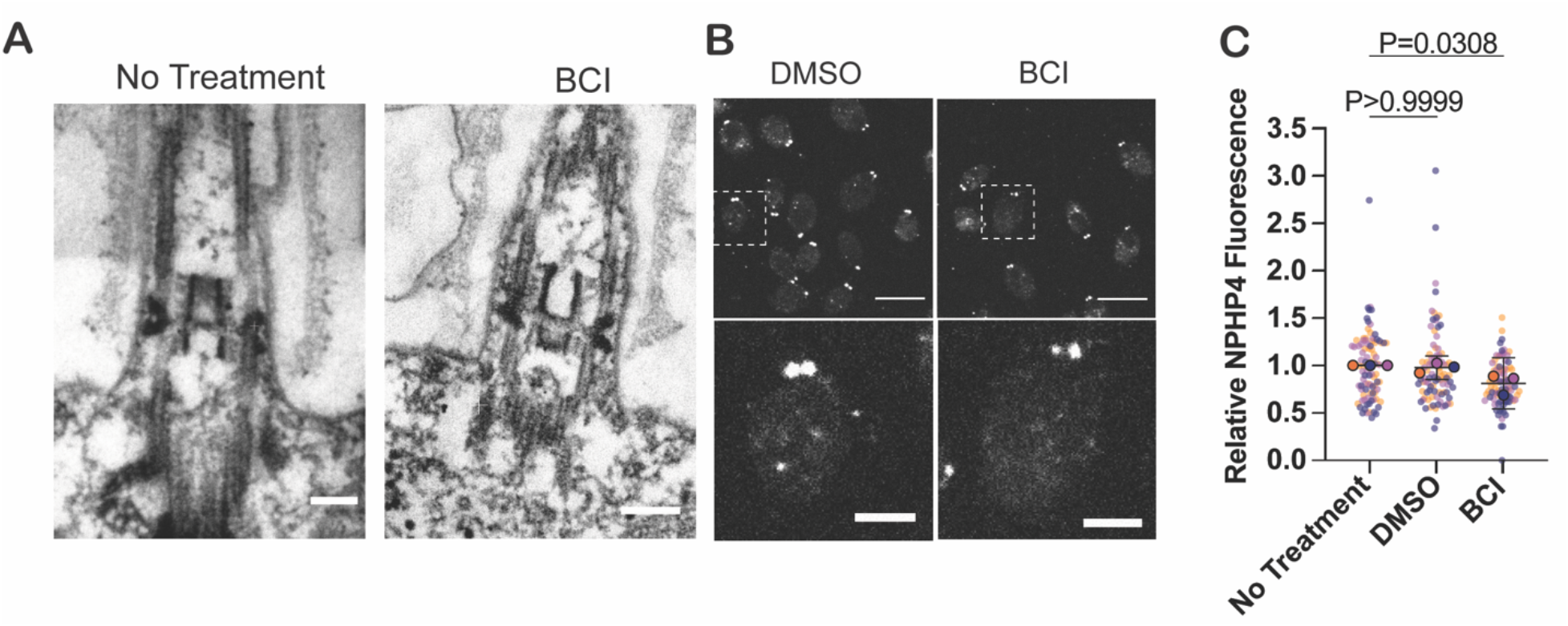
The TZ structure is structurally unaltered with BCI treatment. (**A**) Electron microscopy images of the ciliary transition zone during a 2 hour steady state with or without 30 *µ*M BCI. Scale bars are 100 nm. (**B**) Cells expressing HA tagged NPHP4 were treated with either 0.5% DMSO or 30 *µ*M BCI for two hours and then stained for HA. Scale bars in top images are 5 *µ*m. Scale bars in the bottom insets are 1.5 *µ*m. (**C**) Fluorescent quantification of the NPHP4 signal. Error bars are mean with 95% confidence interval (n=30, N=3). The P values were determined with an ordinary one-way ANOVA with Bonferroni’s multiple comparisons test on the means from each trial.

### BCI disrupts membrane trafficking

The altered KAP-GFP trafficking in the cilium and decreased NPHP4 at the transition zone with a lack of visible structural changes made us curious if other trafficking defects were present in the cell. Membrane trafficking in *Chlamydomonas* has been found to be important for maintaining cilia length and for ciliary assembly. Previous studies have shown that Golgi-derived membrane is important for ciliary assembly and maintenance (Dentler, 2013), and inhibiting internalization through Arp2/3 complex perturbation also results in defective ciliary assembly and maintenance (Bigge et al., 2020). While EM of the Golgi showed no noticeable differences in Golgi structure (**SF 3A**), collapsing the Golgi with BFA (Brefeldin A) intensified ciliary shortening when paired with 20 *µ*M BCI, suggesting that BCI alters other membrane pathways outside of Golgi derived membrane for ciliogenesis (**Fig 5A**). Treatment with 30 *µ*M BCI prevented ciliogenesis post washout, potentially due to toxicity (data not shown).

**Figure 5.**
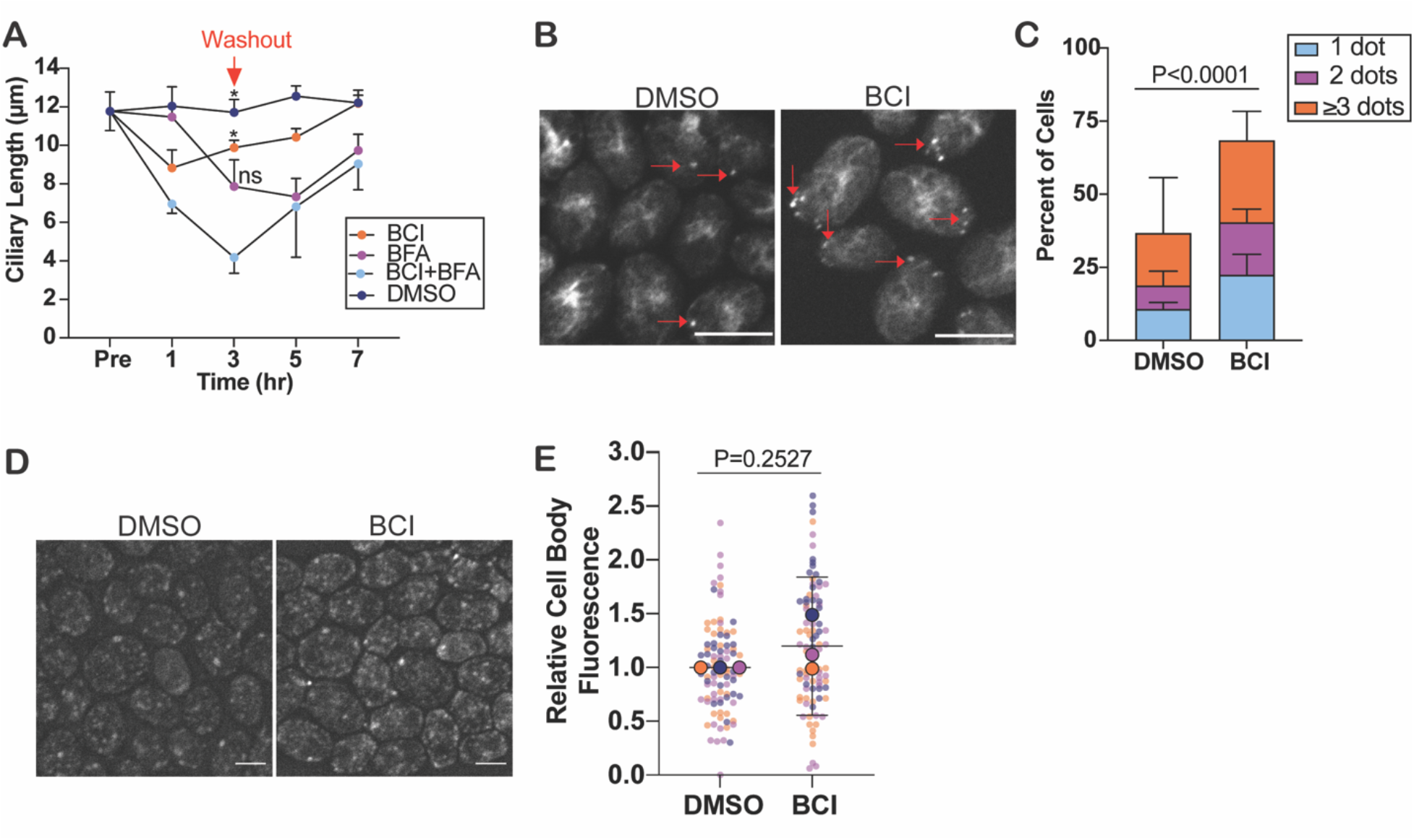
Membrane trafficking is altered in BCI treated cells. **(A)** Steady state cells treated with either 20 *µ*M BCI, 36 *µ*M BFA, both BCI and BFA, or 0.7% DMSO for 3 hours before cells were washed out. Error bars are mean with 95% confidence interval for the mean across 3 experiments (n=30, N=3). The P values compare BCI+BFA to the other treatments at 3 hours and were determined using a two-way ANOVA with Tukey’s correction (* 0.01≤P≤0.05, ns P>0.05). **(B)** Immunofluorescent images of actin puncta visualized with phalloidin in cells treated with 30 *µ*M BCI or DMSO for 2 hours. Red arrows indicate the actin puncta quantified. Scale bars are 5 *µ*m. **(C)** Quantification of actin puncta. Error bars are mean with standard deviation (n=100, N=3). The P value was determined using a Chi-square test with a two-sided Fisher’s exact test for cells with dots versus no dots. **(D)** Immunofluorescent images of membrane dye uptake. Scale bars are 5 *µ*m. **(E)** Quantification of membrane dye uptake. Error bars are mean with the 95% confidence interval (n=30, N=3). The P value was determined with an unpaired t-test.

Previously, Bigge et al. 2020 showed that actin puncta formation, reminiscent of endocytic pits seen in yeast, were absent in ARPC4 cells. In the presence of BCI, actin puncta increased both in steady state cells (**Fig 5B-C**) and in regenerating cells (**SF 3B**). In addition, steady state *Chlamydomonas* DUSP6 orthologs had fewer actin puncta compared to wild type cells (**SF 3C**) indicating that BCI may target proteins in this pathway upstream of MKP2, 3, and 5. We measured total internalization with membrane dye uptake and found that in steady state cells, membrane dye uptake may have increased slightly, but did not differ significantly (**Fig 5D-E**), suggesting that BCI does not alter total internalization. We also checked membrane trafficking through venus-tagged Arl6, a GTPase known to regulate membrane trafficking and is recruited to the basal bodies (Wiens et al., 2010). After regenerating these cells, Arl6 fluorescence slightly increased after 1 hour in the presence of BCI (**SF 3C**). Together, this data suggests that BCI disrupts ciliary membrane trafficking independent of the Golgi. Though plasma membrane derived membrane may have problems being incorporated into cilia, total incorporation in the cell is not disrupted.

### BCI disrupts cytoplasmic microtubule reorganization

To see if cell infrastructure is disrupted, we looked at cytoplasmic microtubule organization in steady state and regenerating cells. Microtubule structure was assigned to each cell as either not present, partially formed where microtubules could be seen at the cell periphery and did not extend to the opposite end, or fully formed where a complete cage was visible extending from one end of the cell to the next end (**Fig 6A**). In regenerating cells, there was a dose-dependent effect on microtubule polymerization where higher concentrations of BCI increasingly inhibited microtubule polymerization post regeneration (**SF4A**). In steady state cells when the microtubules are not typically undergoing global reorganization, BCI did not affect microtubule organization until the microtubules were completely resorbed at higher concentrations of BCI (**SF 4B**). However, upon deciliation, the microtubules undergo large-scale reorganization (Wang et al., 2013). We looked at cells regenerating their cilia and found that cells in BCI could not fully reform their microtubule structure in the presence of BCI (**Fig 6B-E**). To see if cells could be forced to reorganize microtubules in the presence of BCI, we used paclitaxel (PTX) which stabilizes microtubules. In the presence of PTX and BCI, cells were able to reorganize their cytoplasmic microtubules in 60 minutes (**Fig 6B-E**). To rule out the possibility that BCI induces tubulin degradation, we used MG132 which inhibits the proteasome, therefore blocking protein degradation. Cytoplasmic microtubule reorganization was not rescued with MG132 suggesting that tubulin is not degraded in the presence of BCI to prevent reorganization post regeneration (**Fig 6B-E**). These data suggest that BCI inhibits the mechanisms and proteins involved in cytoplasmic microtubule reorganization.

**Figure 6.**
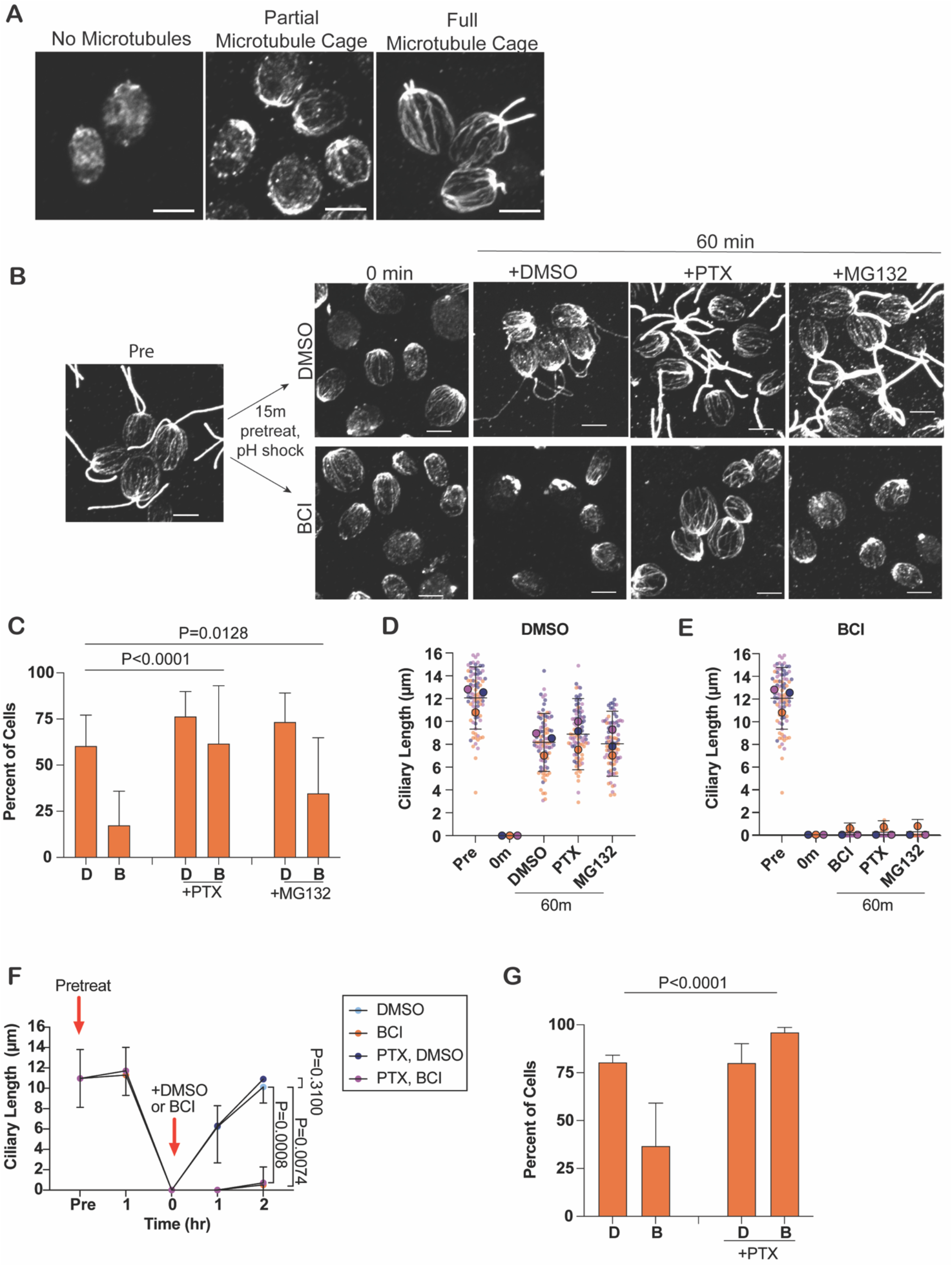
Cytoplasmic microtubule stabilization does not rescue ciliogenesis in the presence of BCI. **(A)** Immunofluorescent examples of the 3 classifications of microtubule structures for microtubule quantification in β-tubulin stained cells: no cage (no microtubules polymerized in the cell), partial recovery (microtubule organization at cell apex, but does not extend to the base), or full recovery (microtubule organization extends fully to both ends of the cell). Scale bars are 5 *µ*m. **(B)** Cells are pretreated with either 0.2% DMSO or 20 *µ*M BCI for 15 minutes and then regenerated in 0.2% DMSO or 20 *µ*M BCI with the addition of 0.5% DMSO (final concentration: 0.7%), 15 *µ*M paclitaxel (PTX), or 50 *µ*M MG132 for 60 minutes before fixing and staining for β-tubulin. Scale bars are 5 *µ*m. **(C)** Quantification of microtubules in (B) at 60 minutes. Bars represent the mean of the percent of cells per category. Error bars are standard deviation (n=100, N=3). D is DMSO, B is BCI. P values were determined using a Chi-square test (p<0.0001) and two-sided Fisher’s exact test. **(D-E)** Quantification of ciliary lengths of cells in (B) pretreated with 0.5% DMSO **(D)** or 20 *µ*M BCI **(E)**. The averages from each trial are plotted on top of the individual data points. Error bars are mean with the 95% confidence interval for the mean from each trial (n=30, N=3). (**F-G**) Cells were pretreated with 0.5% DMSO or 15 *µ*M PTX for 60 minutes and then regenerated with or without Taxol with the addition of either DMSO to 0.7% or 20 *µ*M BCI for 2 hours. **(F)** Ciliary length quantification. Error bars are mean with 95% confidence (n=30, N=3). P values were determined for the 2 hour time points using a two-way ANOVA with Tukey’s correction. **(G)** Quantification of (F) of full B-tubulin reorganization at 120 minutes in 0.7% DMSO or 20 *µ*M BCI with or without 15 *µ*M PTX. Error bars are mean with standard deviation (n=100, N=3). The P value was determined using a Chi-square test (p<0.0001) with a two-sided Fisher’s exact test.

Next, we questioned whether BCI was required for the reorganization of microtubules themselves separate from ciliary assembly. To answer this question, we uncoupled microtubule reorganization from ciliary assembly by treating cells with 2 mg/ml colchicine for 100 minutes to depolymerize cytoplasmic microtubules (Flavin & Slaughter, 1974; LeDizet & Piperno, 1986). Colchicine was then washed out, and microtubules were allowed to reorganize in the presence of DMSO or BCI for 60 minutes and 120 minutes. Interestingly, cells were able to reorganize cytoplasmic microtubules in the presence of BCI, though not to the extent of control cells after 2 hours where ∼50% of control cells had fully repolymerized with 25% of cells only partially reforming cytoplasmic microtubule cages (**SF 4C**). Upon regenerating cells treated for 100 minutes in either Tris-Acetate-Phosphate medium (TAP) or colchicine, those in colchicine could not regenerate cilia nor reorganize their microtubules (**SF 4D**). Together, these data indicate that BCI-induced MAPK activation partially inhibits reorganization of cytoplasmic microtubules during regeneration and slows re-establishment upon colchicine washout in steady state cells.

### Stabilized microtubules do not rescue BCI-defective ciliogenesis

To determine if the defect in microtubule rearrangement after deciliation blocks regeneration in BCI, we stabilized cytoplasmic microtubules with PTX to see if the assembly defect could be rescued. After pretreating cells for 60 minutes with PTX to stabilize the microtubules prior to deciliation and then regenerating in PTX and BCI, ciliogenesis was not able to be rescued, though control cells in PTX were able to regenerate cilia normally (**Fig 6F-G**). This data suggests that though BCI prevents cytoplasmic reorganization, this is not the cause for immediate inhibition to ciliogenesis in regenerating cells. These data are summarized in supplemental figure 5 (**SF 5**). We checked microtubule reorganizaiton in regenerating DUSP6 orthologs and found that none of the ortholog mutants could fully reorganize their microtubules within 60 minutes which suggests a role for these ortholog mutants in regulating cytoplasmic microtubule organization in the cell (**SF 6A-B**). This data further supports that BCI targets proteins involved in cytoplasmic microtubule organization, though this may not directly prevent ciliogenesis from occurring during MAPK activation.

## DISCUSSION

In this study, we have explored mechanisms of action related to ciliogenesis for the DUSP6 inhibitor, BCI, in order to better understand how ERK can regulate ciliogenesis along with the cell cycle. Previous work has shown that silencing the MAPK pathway with MEK1/2 inhibitors can lengthen cilia (S. Wang et al., 2013), and here, we show that through inhibiting the pathway with DUSP6 inhibition, cilia cannot elongate.

We have several possible explanations based on the data to explain why cilia cannot grow in the presence of BCI (**Fig 7**). While kinesin-2 can still be recruited to the basal bodies, its entry is inhibited in regenerating cilia in the presence of BCI, and we have shown that its synthesis is significantly decreased. This data suggests that (1) there could be modifications to existing kinesin-2 in BCI treatment that prevent its entry in ciliogenesis and which slow its kinetics in the cilia and/or (2) the transition zone is altered to prevent kinesin-2 entry. The fact that cilia can grow in cycloheximide to half length suggests that there is enough kinesin-2 and total ciliary protein present at the ciliary base for ciliary incorporation, but BCI immediately prevents incorporation of the precursor pool.

**Figure 7.**
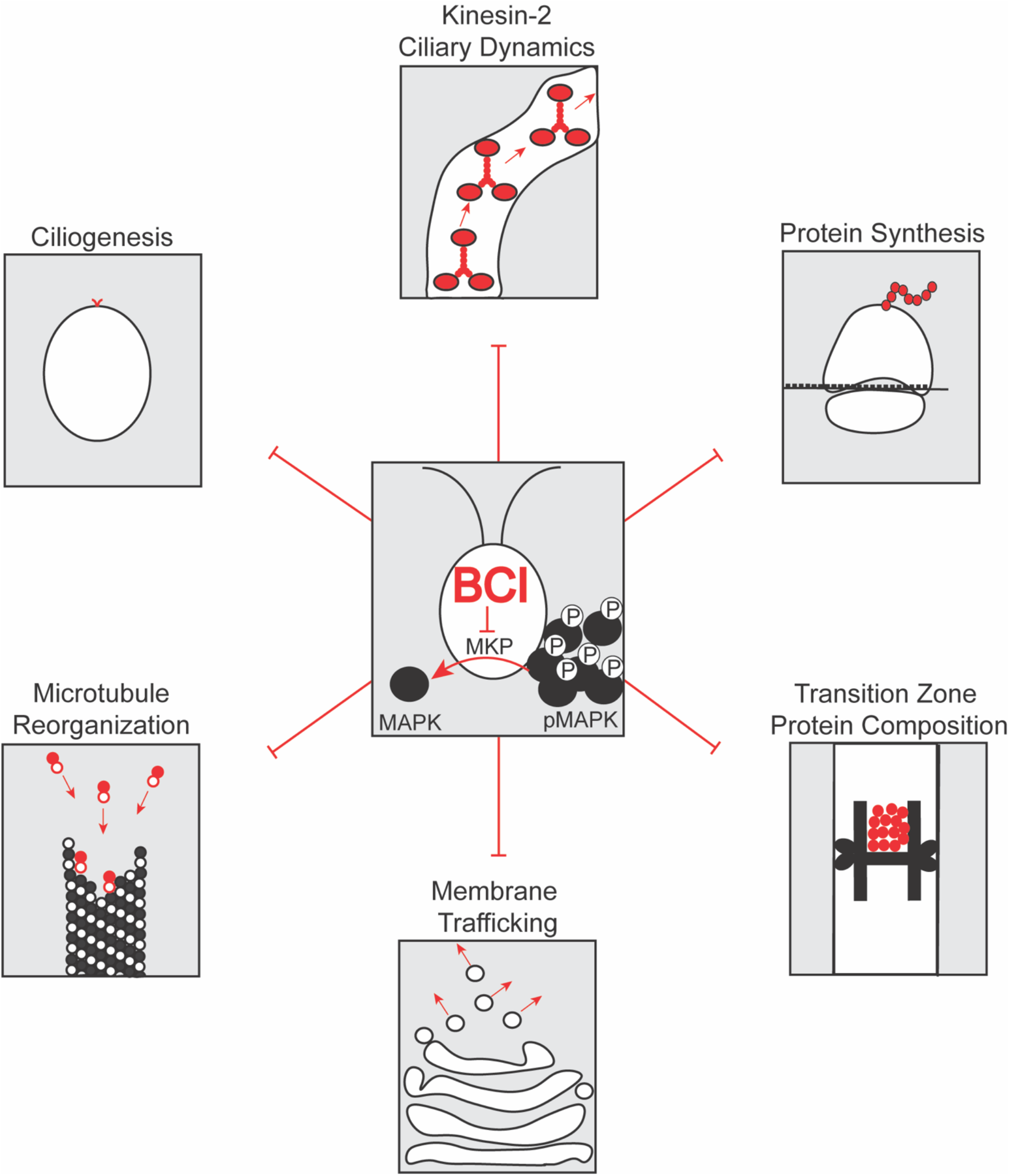
Summary of BCI effects in the cell. BCI inhibits phosphate removal from MAPK. This ultimately alters or inhibits ciliogenesis, kinesin-2 dynamics in cilia, protein synthesis, protein composition at the transition zone, membrane trafficking, and microtubule polymerization.

The transition zone also expresses less NPHP4 in the presence of BCI at a concentration which shortens cilia partially despite the unaltered physical structure (**Fig 4**). The structure could be altered on a finer scale that is not detectable with the techniques employed in this work. The decrease in NPHP4 suggests that there are likely other proteins decreased at the transition zone in a BCI dependent manner such as proteins including NPHP1 with which it directly interacts (Gonçalves & Pelletier, 2017). These proteins may be involved in directly regulating which proteins can enter and exit the cilium, and their absence may slow any movement of proteins into and out of the cilium. It is also possible that MAPK activation results in a phosphorylation cascade which regulates NPHP4 modification, such as through direct phosphorylation, to regulate its localization rather than its protein synthesis. Previous work has shown that phosphorylation of the transition zone protein Tctex1 can promote ciliary disassembly and cell cycle progression (Li et al., 2011). Expression or localization of other proteins in the transition zone may be altered under these conditions.

In addition, we found that there is a membrane trafficking defect. Previous work has shown that inhibition of Golgi derived membrane induces ciliary shortening through Golgi collapse (Dentler, 2013). The epistatic ciliary shortening in BFA and BCI together suggests that BCI may either create an additive effect on the Golgi, or it may disrupt protein and membrane trafficking separate from the Golgi. While the electron microscopy images of the Golgi in BCI did not appear altered, there could be defects that are not visible with this method.

Our data also provides evidence that cytoplasmic microtubule organization does not contribute to early ciliogenesis, though BCI inhibits its reorganization (**Fig 6**). Prevented microtubule polymerization in the presence of BCI following deciliation likely traps kinesin-2 at the basal bodies post deciliation (**Fig 2B**) as well as other proteins required for ciliogenesis at their other locations. Lack of these protein highways could also prevent mRNA from localizing to its intended area for adequate protein synthesis. Constitutive ERK activation could also induce negative feedback loops that prevent transcription as well which would inhibit mRNA formation.

Previous work has provided evidence that microtubule dynamics are required for ciliary assembly in *Chlamydomonas* and destabilization was proposed to be for the purpose of freeing up tubulin for ciliary assembly (Wang et al., 2013). However, microtubule stabilization with paclitaxel before deciliation and the subsequent normal ciliary regrowth suggests that microtubule destabilization is not needed for ciliogenesis (**Fig 6F**). In this study, we used a concentration of paclitaxel that was just low enough to induce stabilized microtubules. It has been shown that lower concentrations of paclitaxel induce G1/G2 arrest whereas higher concentrations induce mitotic arrest (Giannakakou et al., 2001). Similarly, we could be seeing a differential effect for paclitaxel on ciliogenesis where lower concentrations allow ciliogenesis to continue despite the processes induced in addition to microtubule stabilization.

Together, we have found mechanisms contributing to ciliary defects which could be the foundation of various ciliary diseases; the cilium acts as a major signaling hub in the cell to regulate cellular activities. When this structure cannot form and function properly on ciliated cells, important signals cannot be transmitted throughout the cell. Cells normally resorb their cilia for division and then reassemble them during G1/G0. BCI could arrest cells at the interphase checkpoint or induce cell cycle arrest which could inhibit protein synthesis to shift ciliary assembly to disassembly. There is normally constant turnover of tubulin and ciliary proteins which is normally replaced and because BCI stops protein synthesis (**Fig 3J**), cilia do not have the supplies to keep growing cilia and thus we see ciliary shortening (Song & Dentler, 2001; Stephens, 1997). While the protein synthesis defect does not explain why there is complete and immediate inhibition of ciliogenesis in regenerating cells, it can help explain why cilia cannot grow over time with MAPK activation.

## MATERIALS AND METHODS

### Strains and Maintenance

The wild type strain (CC-5325), NPHP4-HAC, KAP-GFP, *mkp2* mutant (LMJ.RY0402.168348), *mkp3* mutant (LMJ.RY0402.191934), and *mkp5* mutant (LMJ.RY0402.083264), and Arl6 strain were acquired from the *Chlamydomonas* Resource Center. *Chlamydomonas* cells were grown on 1.5% Tris-Acetate-Phosphate (TAP) agar plates in constant light (Helloify BR30 LED Plant Grow Light Bulb, 9W, 14umol/s blue and red light). Cells were inoculated in liquid TAP and grown in constant light and agitation 18 hours prior to experiments.

The human TERT-RPE1 cells were a gift from Chris Shoemaker at Dartmouth College, Hanover, NH. These cells were maintained in Dulbecco’s Modified Eagle Medium (DMEM, Corning MT10013CV) with 10% fetal bovine serum (FBS) and 10 µg/mL Hygromycin B (InvivoGen) at 37°C in 5% CO_2_. To induce ciliogenesis, cells were grown to 80% confluency and then serum starved in DMEM with 0.5% FBS for 48 hours at 37°C, 5% CO_2_ prior to experiments.

### Genotyping

A single colony of *Chlamydomonas* cells were placed in chelex beads, boiled at 90°C, and then spun down at 550xg, 1min. This template was used along with 1x DreamTaq Buffer (Invitrogen), 2 mM dNTPs (NEB), 0.5 *µ*M forward and reverse primers (IDT), and Dream Taq Polymerase (Invitrogen). PCR parameters were used according to the *Chlamydomonas* Resource Center. DNA was mixed with 6x loading dye to 1x (Invitrogen) and run on a 1% agarose gel in TBE with 1x SYBR Safe DNA Gel Stain (Invitrogen, S33102) and imaged on a Bio-Rad ChemiDoc MP.

### Primers

*mkp2* mutant forward primer : CTGCGGACATCAGCTCAAT

*mkp3* mutant forward primer: CAAGAGCACCTGGCACAGGAG

*mkp5* mutant forward primer: TCGTGACAGACCTGCAGAG

CIB1 reverse primer: CCGAGGAGAAACTGGCCTTCT

### Ciliary Length Experiments

Steady state cells were grown in liquid TAP under constant light and agitation for 18 hours and then treated with >1% DMSO or BCI (Sigma, B4313-5MG) for 2 hours. Samples were collected in equal volumes of 2% glutaraldehyde and allowed to settle before imaging. To measure cilia, 3uL of settled cells were placed directly on a slide with a coverslip and imaged at 40x magnification with a Zeiss Axioscope 5 DIC and Zeiss Zen 3.1 software. One cilium per cell was measured with the segmented line tool and fit spline for 30 cells per time point.

Regenerating cells were grown in liquid TAP under constant light and agitation for 18 hours and then pH shocked for 45 seconds with 0.5M acetic acid to bring the culture pH=4.5 and then brought back up to pH=7.0 with 0.5M potassium hydroxide. These cells were immediately centrifuged at 550xg for 1 minute and supplied new TAP with or without drugs. For double regenerations, cells were allowed to grow for 1 hour in 10 *µ*g/mL cycloheximide (Sigma, C1988-1G) and then pH shocked a second time, resuspended in new TAP, and grown with or without drugs. Samples for DIC imaging were collected in equal volumes of 2% glutaraldehyde.

### TIRF Microscopy and Quantification

Cells were treated with 0.5% DMSO or 30 *µ*M BCI for 2 hours, staggered to allow time for imaging at both 1 hour and 2 hours of treatment. KAP-GFP cells were placed on a poly-lysine treated coverslip and diluted 1:100 in TAP. Cells were imaged with a 100x oil immersion objective on a Nikon Eclipse Ti microscope with two Andor 897 EMCCD cameras suing the 499nm laser line. Data was collected and kymographs generated in NIS Elements Software.

Quantifications were carried out on the kymographs in FIJI. KAP-GFP trains (groups of KAP-GFP molecules) are indicated by the fluorescing slants in the kymograph. For frequency measurements Frequency = (number of trains)*(Frames per second/total frames measured). For velocity: Velocity = (tanθ in radians of train relative to the basal body)*(Frames per second)*(um/pixel conversion). For injection size (fluorescent intensity of a train), kymographs were background subtracted with rolling ball radius and a line thick enough to cover the largest GFP signal was drawn over each train from the basal bodies to the tip of the cilium and then normalized. Measurements were repeated in Kymograph Direct (Mangeol et al., 2016) (data not shown) to ensure similar results were obtained.

### Cilia Isolation

Cilia were isolated according to Craige et al., 2013. Cells were grown in TAP with constant light and agitation. Before isolating cilia, cells were treated with 30 *µ*M BCI or 0.5% DMSO for 2 hours. Cells were spun down at ∼550xg for 5 minutes at room temp and resuspended in 10mM HEPES (pH=7.4). Cells were spun down at 550xg for 5 minutes and resuspended in cold 4% HMDS (10mM HEPES, PH=7.4, 5mM MgSO4, 1mM DTT, and 4% (w/v) sucrose) and then incubated with 25 mM dibucaine for 2 minutes before adding HMDS with 0.5mM EGTA. Cilia were separated from the cell bodies by centrifuging for 5 minutes at 1800 rpm and then purified with a sucrose gradient by underlaying 25% sucrose in HMDS. The gradient was spun for 10 minutes, 2400xg, 4°C without braking. The top half of the gradient was further spun at 21,000xg for 30 minutes, 4°C and the pellets were collected in small amounts of lysis buffer for western blotting (Craige et al., 2013).

### Immunofluorescence and Quantification

#### KAP-GFP immunofluorescence and quantification

Cells were adhered to poly-lysine coated coverslips for 2 minutes and then permeabilized in cold (−20°C) methanol for 10 minutes, replacing the methanol once after 5 minutes. Coverslips were allowed to dry and then mounted with Fluoromount G (Thermo Scientific) and imaged with 100x oil immersion objective on a spinning disk confocal microscope (Nikon Eclipse Ti-E with Yokogawa 2-camera CSU0W1 spinning disk system and Nikon LU-N4 laser launch). To quantify basal body fluorescence and cilia fluorescence, summed slice projections were collected using FIJI for each image stack and background subtracted with rolling ball radius. An equal size circle was drawn around the basal bodies across one trial and the area, mean gray value, and integrated density measurements were collected for 10 sets of basal bodies for each of 3 pictures per time point or treatment group. The total cell fluorescence was calculated using the calculation for corrected total cell fluorescence (CTCF=IntDen-([Area of selection]*[background mean grey value]). Background mean grey values were calculated by taking the average of 3 random background measurements per image. For cilia measurements, the selection area involved drawing a tight area around the cilia for one cilium per cell, 10 cells per image, and 3 images per time point or treatment. The length of the cilia was measured alongside the fluorescence measurements and plotted against each other (**Fig 2C**).

#### Microtubule immunofluorescence and quantification

Staining was performed similarly to Wang et. al, 2013. Cells were adhered to poly-lysine coverslips, fixed in microtubule buffer (30 mM HEPES [pH=7.2], 3 mM EGTA, 1 mM MgSO_4_, 25 mM KCl) containing fresh (<1 month opened) 4% PFA for 5 minutes, incubated in microtubule buffer with 0.5% NP-40 for 5 minutes, and then permeabilized in cold (−20°C) methanol for 5 minutes. Coverslips were placed in a humidified chamber and blocked in 5% BSA and 1% fish gelatin for 30 minutes and 10% normal goat serum in block for 30 minutes. Coverslips were then incubated overnight at 4°C with primary antibody (B-tubulin, CST 2146S, 1:100) in 20% block in 1x PBS. Washes were done with 1x PBS, 3×10 minutes and coverslips were placed back in humidified chamber with secondary antibody (Alexa Fluor 488 goat anti-rabbit IgG, Invitrogen A11008, 1:500) for 1 hour covered. Coverslips were washed 3×10 minutes covered and allowed to dry completely before mounting with Fluoromount G and imaging on a spinning disk confocal microscope. Microtubule quantification was performed as described in Figure 6 where 100 cells total were counted and assigned to have either no microtubules, partially visible microtubules, or full microtubules according to the staining. Cells were counted using the cell counter plugin in FIJI.

#### Phalloidin Staining

Phalloidin staining was performed according to Craig and Avasthi, 2019. Cells adhered to coverslips for 2 minutes and were fixed with fresh 4% PFA in 7.5 mM HEPES in 1x PBS for 15 minutes, washed in 1x PBS for 3 minutes, permeabilized in cold (−20°C) 80% acetone for 5 minutes and then 100% cold acetone and allowed to air dry. Coverslips were rehydrated in 1x PBS for 5 minutes and incubated with Atto-488 Phalloidin (Sigma, 49409-10Nmol) for 16 minutes and then washed in 1x PBS for 5 minutes, covered (Craig & Avasthi, 2019). Coverslips were air dried, mounted in Fluoromount G, and imaged at 100x on the spinning disk confocal microscope. For dot quantification, max intensity projections were generated for each image stack and dots per cell counted using FIJI’s cell counter plugin.

#### Membrane Staining

Membrane staining was performed according to Bigge et. al, 2020. Cells were treated with 0.5% DMSO or 30 *µ*M BCI for 2 hours and then adhered to coverslips for 2 minutes. Coverslips were moved to ice and 200 *µ*g/mL membrane dye (FMTM 4-64FX, fixable; Thermo, F34653) for 1 minute covered, fixed with 4% PFA in 1x HBSS for 10 minutes, and washed 3×3 minutes in 1x PBS (Bigge et al., 2020). Coverslips were air dried and mounted to slides with Fluoromount G and imaged on the spinning disk confocal microscope. For quantification, summed slice projections were created and processed as described in KAP-GFP staining. The area selection measured encompassed an entire cell for 10 cells per image, 3 images per treatment and time point, across 3 trials.

#### RPE1 cilium staining

Staining was performed according to Alsolami et al., 2019. Cells were seeded on chamber slides and grown to 80% confluency before serum starving in DMEM with 0.5% FBS for 48 hours. Media was removed and cells were washed with 1x PBS, fixed in cold (−20°C) methanol for 10 minutes and washed once more in 1x PBS for 5 minutes. Cells were blocked in 1x PBS with 0.2% Triton X-100 (PBX) at 37°C for 5 minutes and then in PBX with 3% BSA for 30 minutes at 37°C. Slides were moved to a humidified chamber with primary antibody (anti-acetyl-alpha tubulin, clone 6-11B-1, Sigma, MABT868) diluted in PBX (1:250) added for 1 hour at room temp. Slides were washed 3×5 minutes in PBS and placed back in a covered humidified chamber. Secondary antibody (Alexa Fluor 488 goat anti-mouse IgG, Invitrogen, A11001) was diluted in PBX (1:500) and added to coverslips for 1 hour at room temp. Slides were washed 3×5 minutes in PBS, mounted with Flouromount G, and imaged with 60x and 100x oil immersion objectives on the spinning disk confocal (Alsolami et al., 2019). Max intensity projections were created in FIJI for each image, and primary cilia were measured using the segmented line tool for the acetylated tubulin staining.

#### Sytox Green and Hoechst Staining

Cells were seeded in chamber slides and grown to 80% confluency before serum starving in DMEM with 0.5% FBS for 48 hours. Media was removed from RPE1 cells and replaced with warmed 1x HBSS containing 1:500 dilutions of both Sytox Green nucleic acid stain (Life Technologies, S7020, 1:500 dilution) and Hoechst (Thermo, 62249) for 30 minutes at 37°C, covered. Slides were washed 3×5 minutes in HBSS. A coverslip was added to the slides directly and cells were imaged with a 60x immersion oil objective on the spinning disk confocal. For quantification, max intensity projections were generated and 100 cells were counted total per concentration. Sytox green nuclear staining indicated dead cells whereas Hoechst stained nuclei in total cells.

### SDS-PAGE and Immunoblotting

RPE1 cells were washed with cold 1x HBSS and then placed on ice with RIPA buffer + 10x phosphatase inhibitors for 20 minutes with agitation after 10 minutes. Cells were scraped and placed into a tube and spun at 21,000xg, 10min, 4°C. The lysate was mixed with 4x loading buffer (Invitrogen) and BME to 10%, boiled at 95°C for 10 minutes, and run on a 10% bis-tris gel for 2 hours before transferring to a PVDF membrane and blocked for 1 hour in TBS-T and 5% BSA. The primary antibody (Phospho p44/42 MAPK (Erk1/2) (Thr202/Tyr204) Antibody #9101, CST) was diluted 1:1000 in TBS-T and incubated with blots overnight at 4°C. Blots were washed 3 times in TBS-T and incubated with secondary antibody (Goat anti-rabbit IgG HRP conjugate, Invitrogen, G21234) for 1 hour at room temp. Blots were washed 3×10 minutes in TBS-T and then incubated with pico chemiluminescent substrate (Invitrogen) and imaged on a Bio-Rad ChemiDoc MP.

*Chlamydomonas* cells were centrifuged at 550xg for 1 minute. The supernatant was removed and replaced with 100uL of glass beads (425-600 *µ*m) and 100 *µ*L lysis buffer (1% NP-40, 9% TAP, 5% glycerol, 1mM DTT, 1x protease and phosphatase inhibitors per 1mL of cells spun down. Cells were bead beated 3×1minute with 1 minute breaks on ice and then centrifuged at 21,000xg for 15 minutes at 4°C. The supernatant was collected and mixed with DTT to a final concentration of (50 *µ*M?) and to 1x LDS Sample Buffer (Invitrogen), and boiled at 70°C for 10 minutes. Protein was run on 10% Tris-bis gels (Invitrogen) at 150V and transferred onto PVDF membrane.

To probe for KAP-GFP, blots were incubated with 5% milk and then probed with GFP antibody (CST, 1956S) in 1% milk +BSA overnight at 4°C. Blots were washed 3×10 minutes in PBS-T and incubated with secondary antibody (goat anti-rabbit IgG HRP conjugate, Invitrogen, G21234) diluted in 1% milk +BSA (1:5000) for 1 hour at room temp. Blots were washed 3×10 minutes in TBS-T and then incubated with pico chemiluminescent substrate (Thermo) and imaged on a Bio-Rad ChemiDoc MP.

### Electron Microscopy

Electron microscopy was performed by Radu Stan according to a protocol by Dentler and Adams 1992. Wild type cells were treated with 30 *µ*M BCI or 0.5% DMSO for 2 hours and then diluted in equal volumes of fresh, EM-grade 2% glutaraldehyde diluted from 16% in water (Electron Microscopy Sciences). Cells settled at room temperature and the supernatant was removed and cells were resuspended with 1% glutaraldehyde with 50 mM sodium cacodylate overnight at 4°C. The supernatant was removed and cells were spun 3×1 minute at 600xg with 50 mM sodium cacodylate washes and then fixed and imaged with a Helios Scanning Electron Microscope 5CX with a STEM3+ detector (Dentler & Adams, 1992).

### Statistical Analysis

Data collection and statistical calculations were completed in GraphPad Prism Version 9.2.0 (283). Super plots were generated according to (Lord et al., 2020). In figure legends, n refers to individual measurements made across one trial, and N refers to the number of trials performed. Geneious Prime 2022 was used to generate the phylogenetic tree and sequence alignment (**SF1A-B**). *Chlamydomonas* sequences were generated from Phytozome 13 (Goodstein et al., 2012).

## ACKNOWLEDGEMENTS

We are extremely grateful for Chris Shoemaker and his lab for providing hTERT-RPE1 cells, methods, and guidance on related procedures to facilitate this work. We would also like to thank Ann Lavanway at the Dartmouth Imaging Facility for helping with immunofluorescence imaging, Radu Stan for preparing and imaging cells for electron microscopy (funding source?), the BioMT Core at Dartmouth (funded by NIH NIGMS grant P20-GM113132) and Genomics and Molecular Biology Shared Resources Core at Dartmouth (funded by NCI Cancer Center Support grant 5P30CA023108-37) for use of equipment, the Department of Anatomy and Cell Biology and the University of Kansas Medical Center where this work was started, and our funding source NIH MIRA R35GM128702(PA).

## SUPPLEMENTAL FIGURES

**Supplemental Figure 1.**
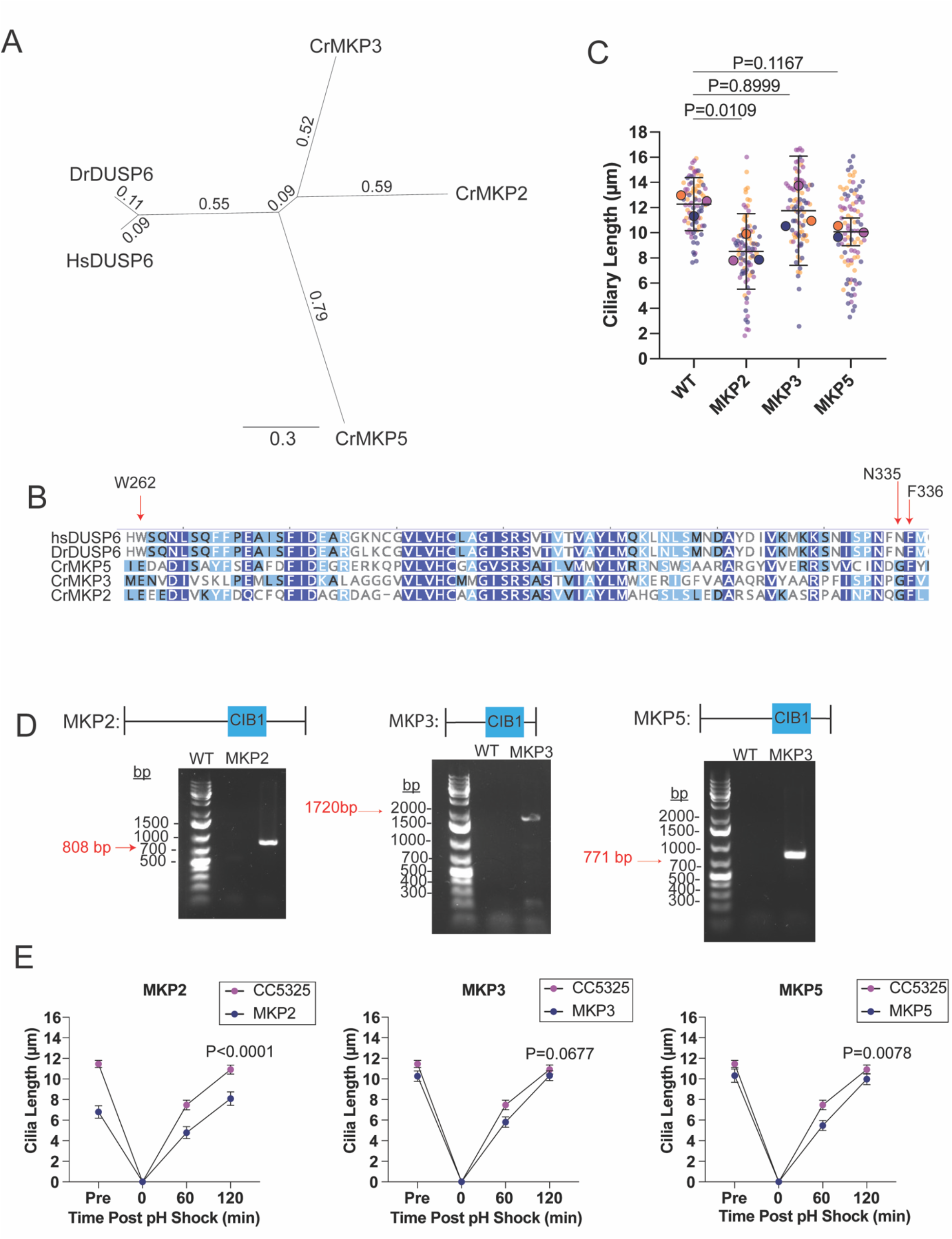
*Chlamydomonas* DUSP6 orthologs have ciliary length defects. (**A)** Phylogenetic tree for the top 3 zebrafish DUSP6 hits in *Chlamydomonas*. The phylogenetic tree is a neighbor-joining tree. Numbers denote branch length. **(B)** Sequence alignment of predicted residues BCI interacts with (Molina et al., 2009). **(C)** Genotype results for the *Chlamydomonas* DUSP6 orthologs acquired from the *Chlamydomonas* Resource Center. **(D)** Steady state lengths of the top 3 most similar *Chlamydomonas* mutants. Error bars are mean with 95% confidence interval (n=30, N=3). P values were determined using an ordinary one-way ANOVA with Dunnett’s correction. **(E)** The mean ciliary length measurements with mean and 95% confidence intervals in regenerating DUSP6 mutants over 2 hours (n=30, N=3). P values compare 120 minute time points in each case which were determine with an unpaired t-test.

**Supplemental Figure 2.**
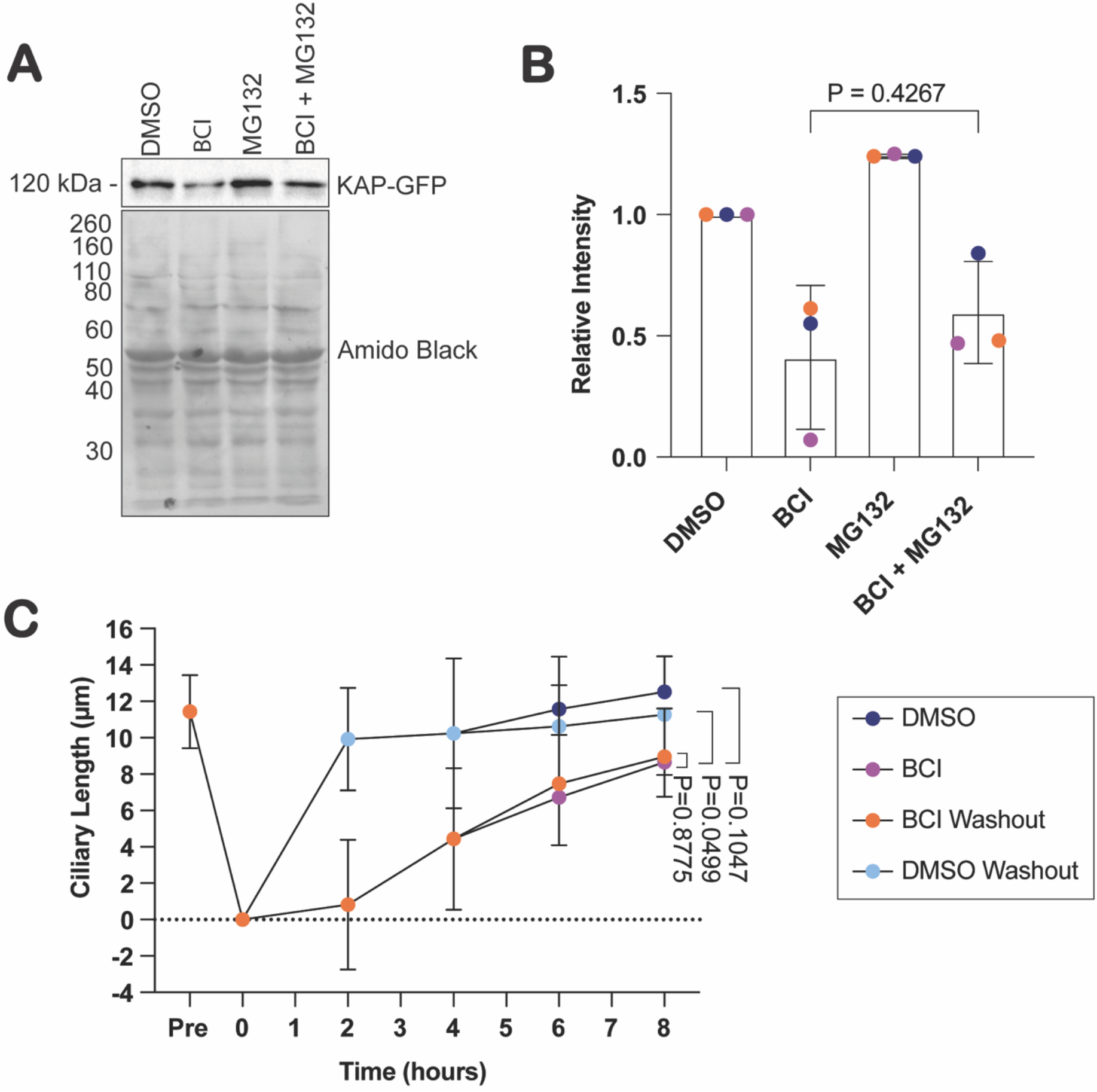
BCI inhibits protein synthesis. **(A**) Western blot of KAP-GFP expression compared to total protein exposed with amido black. Cells were treated for 2 hours with either 0.5% DMSO, 30 *µ*M BCI, 50 *µ*M MG132, or both 50 *µ*M MG132 and 30 *µ*MBCI. (**B**) Quantification of (A). Error bars are standard deviation of the mean (n=1, N=3). The P value was determined by an unpaired t-test between BCI and BCI+MG132. **(C)** Single regeneration performed in parallel with Figure 3H. Error bars are mean with 95% confidence interval (n=30, N=3). P values were determined with a two-way ANOVA with Tukey’s correction for multiple comparisons.

**Supplemental Figure 3.**
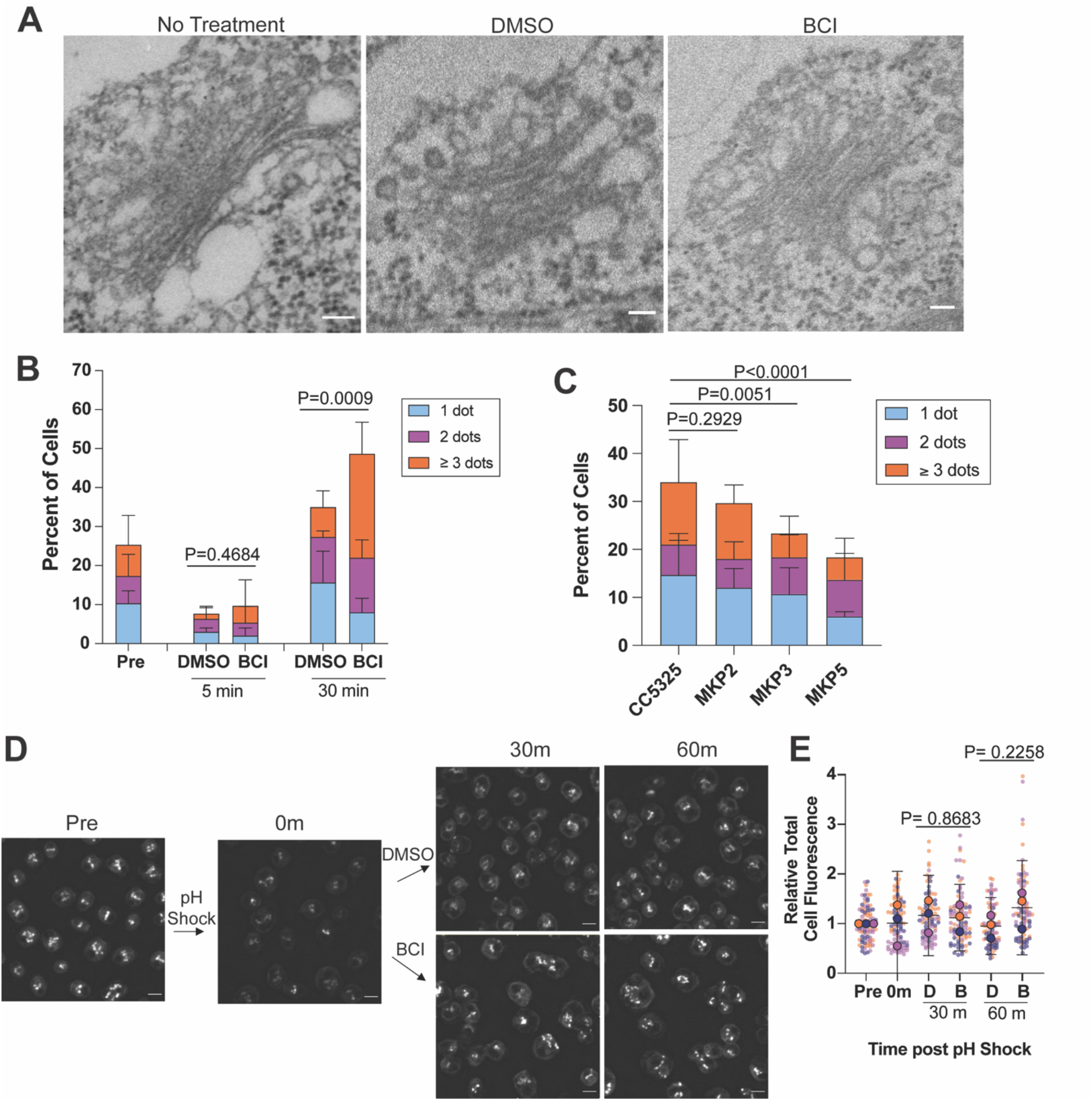
*Chlamydomonas* DUSP6 orthologs and tagged Arl6 show membrane trafficking defects. (**A**) Representative EM images of the Golgi apparatus in cells with no treatment, 0.5% DMSO, or 30 *µ*M BCI. Scale bars are 100nm. (**B**) Quantification of actin puncta in regenerating wild type cells in either BCI or DMSO. Error bars are mean with standard deviation (n=100, N=3). P values were determined using an unpaired t-test. (**C**) Quantification of actin puncta in steady state DUSP6 orthologs. Error bars are mean with standard deviation (n=100, N=3). P values were determined using a Chi-square test (P<0.0001) and a two-sided Fisher’s exact test comparing no dots to total dots. Individual bars represent average percent of cells (n=100, N=3). (**D**) Representative images of Arl6/F02 venus-tagged *Chlamydomonas* cells regenerated in 0.5% DMSO or 30 *µ*M BCI for 1 hour. (**E**) Quantification of arl6 whole cell fluorescence. Error bars are mean with 95% confidence interval (n=30, N=3). P values were determined using an unpaired t test.

**Supplemental Figure 4.**
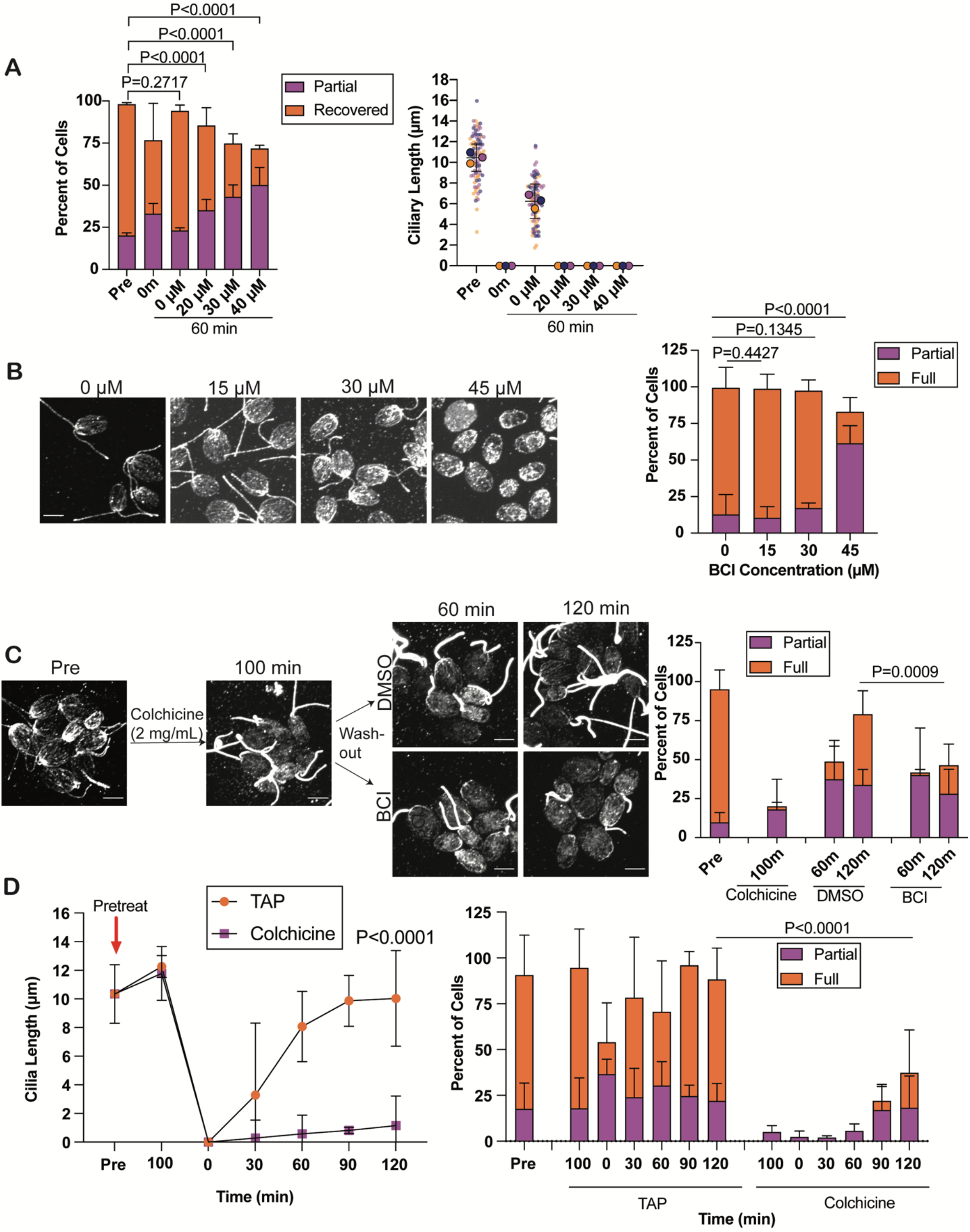
Initial colchicine-induced depolymerization prevents ciliogenesis and microtubule reorganization. (**A**) Cells were regenerated in either 0.4% DMSO or increasing concentrations of BCI and stained for B-tubulin. Error bars are the mean with standard deviation for the microtubule quantification (left) (n=100, N=3). P values for microtubule reorganizaiton were determined using a Chi-square test with a two-sided Fisher’s exact test. The ciliary length quantification (right) shows the mean with the 95% confidence interval (n=30, N=3). (**B**) Steady state wild type cells were treated with either 0.5% DMSO or increasing concentrations of BCI for 2 hours and stained for B-tubulin. Error bars are mean with standard deviation (n=100, N=3). P values were determined with a Chi-square test (p<0.0001) and a two-sided Fisher’s exact test. (**C**) Cells were pretreated with 2 mg/mL colchicine for 100 minutes. Colchicine was washed out and cells were incubated with either 0.2% DMSO or 20 *µ*M BCI immediately following. Cells were stained for B-tubulin. Microtubule repolymerization is quantified on the right. Error bars are mean with standard deviation (n=100, N=3). P values were determined using a Chi-square test (p=0.0009) and two-sided Fisher’s exact test for the 120 minute time points. (**D**) Cells were pretreated with either TAP or 2 mg/mL colchicine for 100 minutes and then regenerated in TAP. Cells were fixed and stained for B-tubulin. For ciliary length measurements, error bars are mean with 95% confidence interval (n=30, N=3). The P value for the 120 minute ciliary timepoints was determined using a two-way ANOVA with Tukey’s correction. Microtubule repolymerization is quantified on the right. Error bars are mean with standard deviation (n=100, N=3). P values for the 120 minute microtubule reorganization were determined using a Chi-square test (p=0.0008) and two-sided Fisher’s exact test.

**Supplemental Figure 5.**
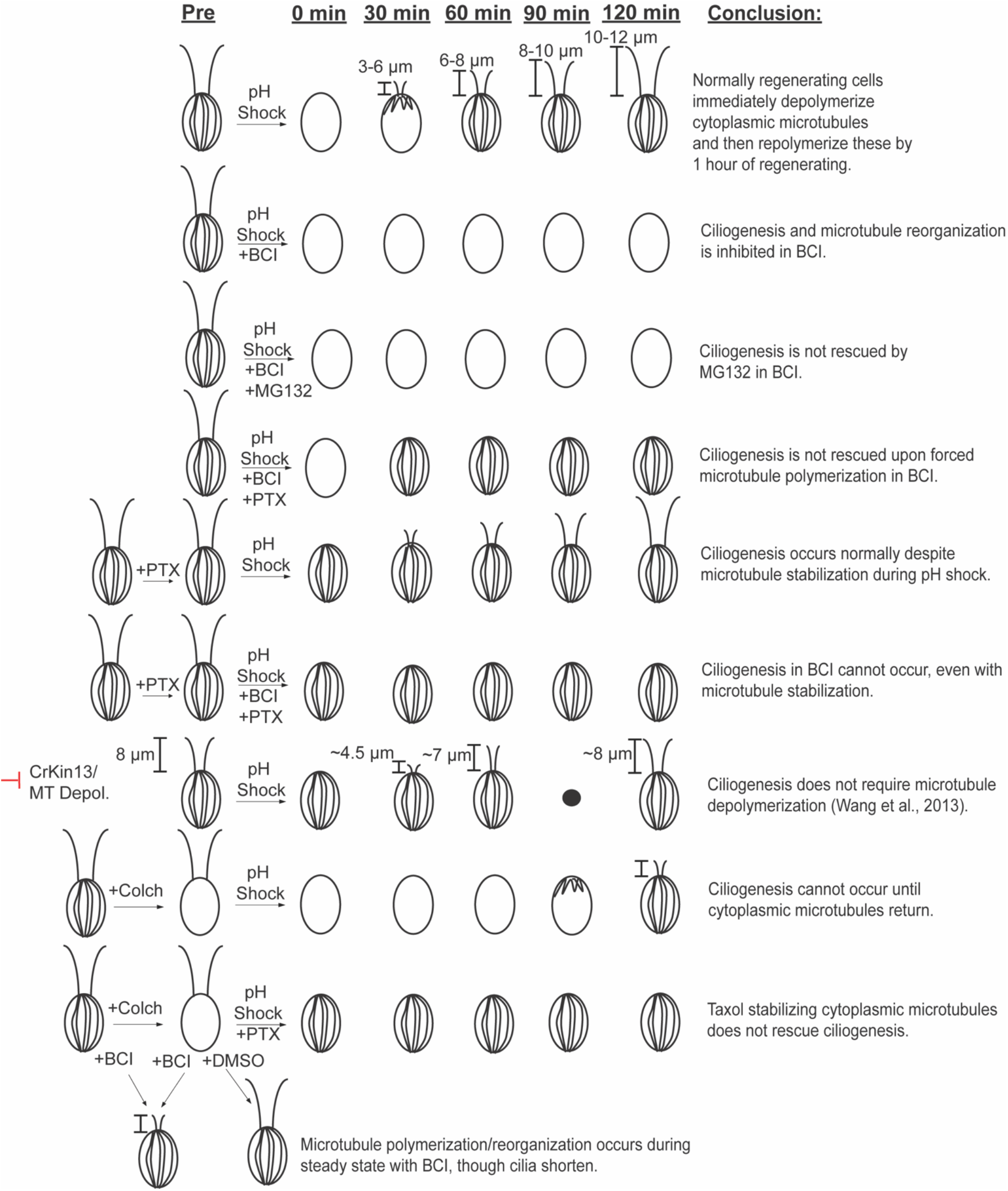
Summary of microtubule data.

**Supplemental Figure 6.**
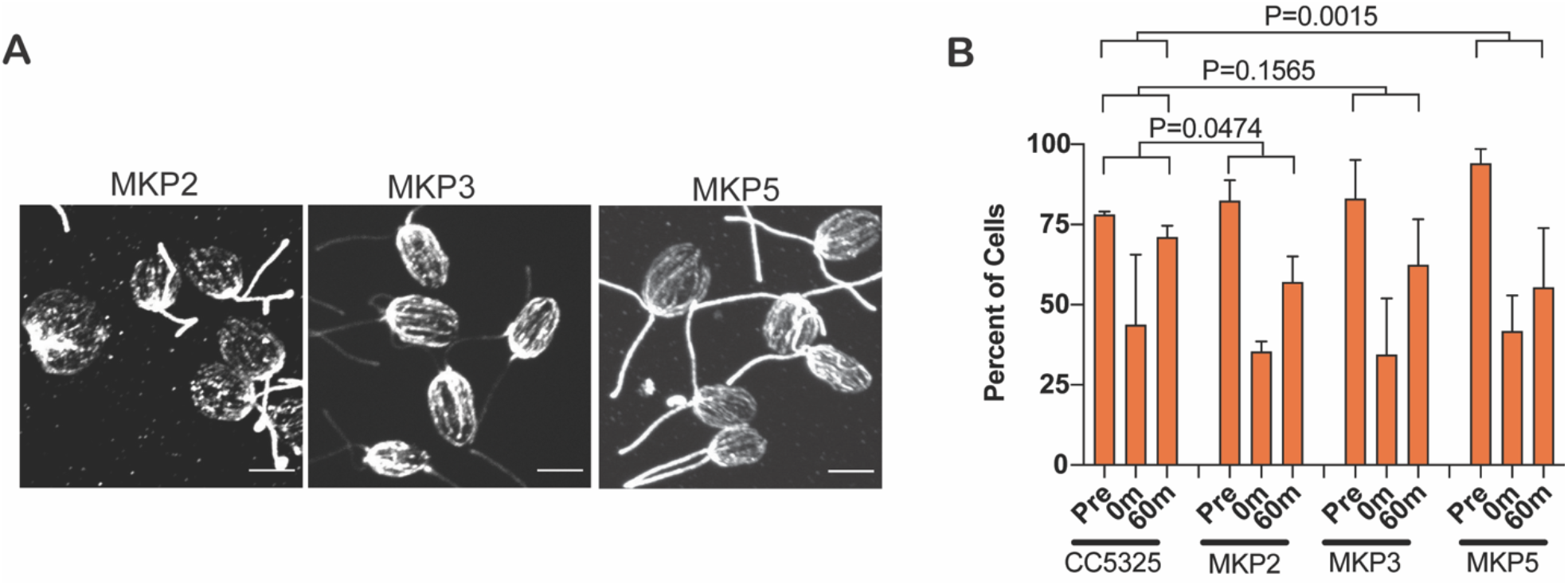
*Chlamydomonas* DUSP6 orthologs exhibit microtubule polymerization defects. (**A**) *Chlamydomonas* DUSP6 orthologs were regenerated and stained for B-tubulin. Scale bars are 5 *µ*m. (**B**) The mean number of cells over 3 trials is plotted with standard deviation (n=100, N=3). P values compare the pre and 60 minute full microtubule quantifications between CC5325 and each mutant. The P values were calculated using a Chi-square test (P=0.0136) with a two-sided Fisher’s exact test.

## Notes

### Competing Interest Statement

PA is a paid consultant for Arcadia Science

