## Supplemental Material for "ERK pathway activation inhibits ciliogenesis and causes defects in motor behavior, ciliary gating, and cytoskeletal rearrangement"

### SUPPLEMENTAL FIGURES

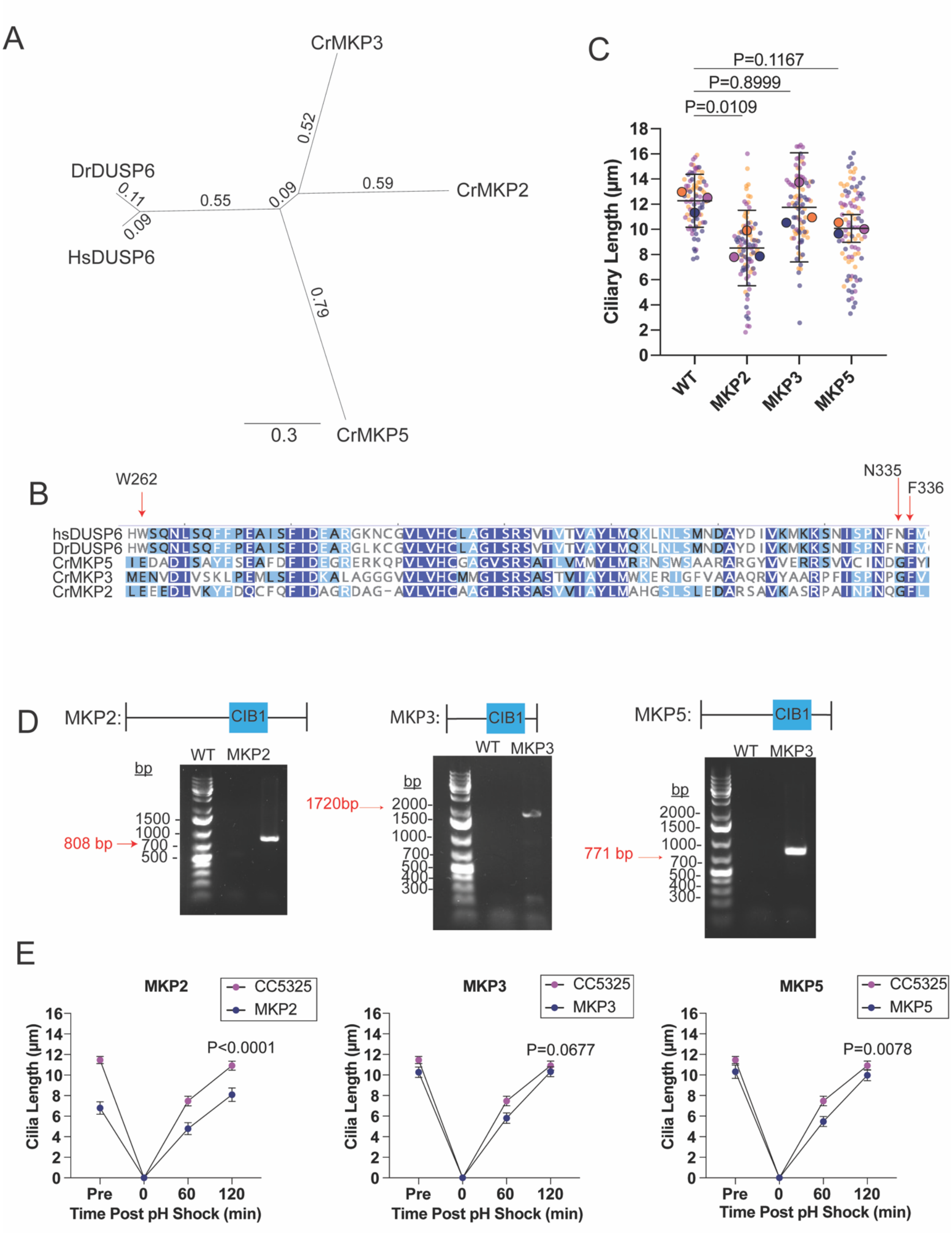

**Supplemental Figure 1.** ***Chlamydomonas*** **DUSP6 orthologs have ciliary length defects**. (**A**) Phylogenetic tree for the top 3 zebrafish DUSP6 hits in *Chlamydomonas*. The phylogenetic tree is a neighbor-joining tree. Numbers denote branch length. **(B)** Sequence alignment of predicted residues BCI interacts with (Molina et al., 2009). **(C)** Genotype results for the *Chlamydomonas* DUSP6 orthologs acquired from the *Chlamydomonas* Resource Center. **(D)** Steady state lengths of the top 3 most similar *Chlamydomonas* mutants. Error bars are mean with 95% confidence interval (n=30, N=3). P values were determined using an ordinary one-way ANOVA with Dunnett's correction. **(E)** The mean ciliary length measurements with mean and 95% confidence intervals in regenerating DUSP6 mutants over 2 hours (n=30, N=3). P values compare 120 minute time points in each case which were determine with an unpaired t-test.

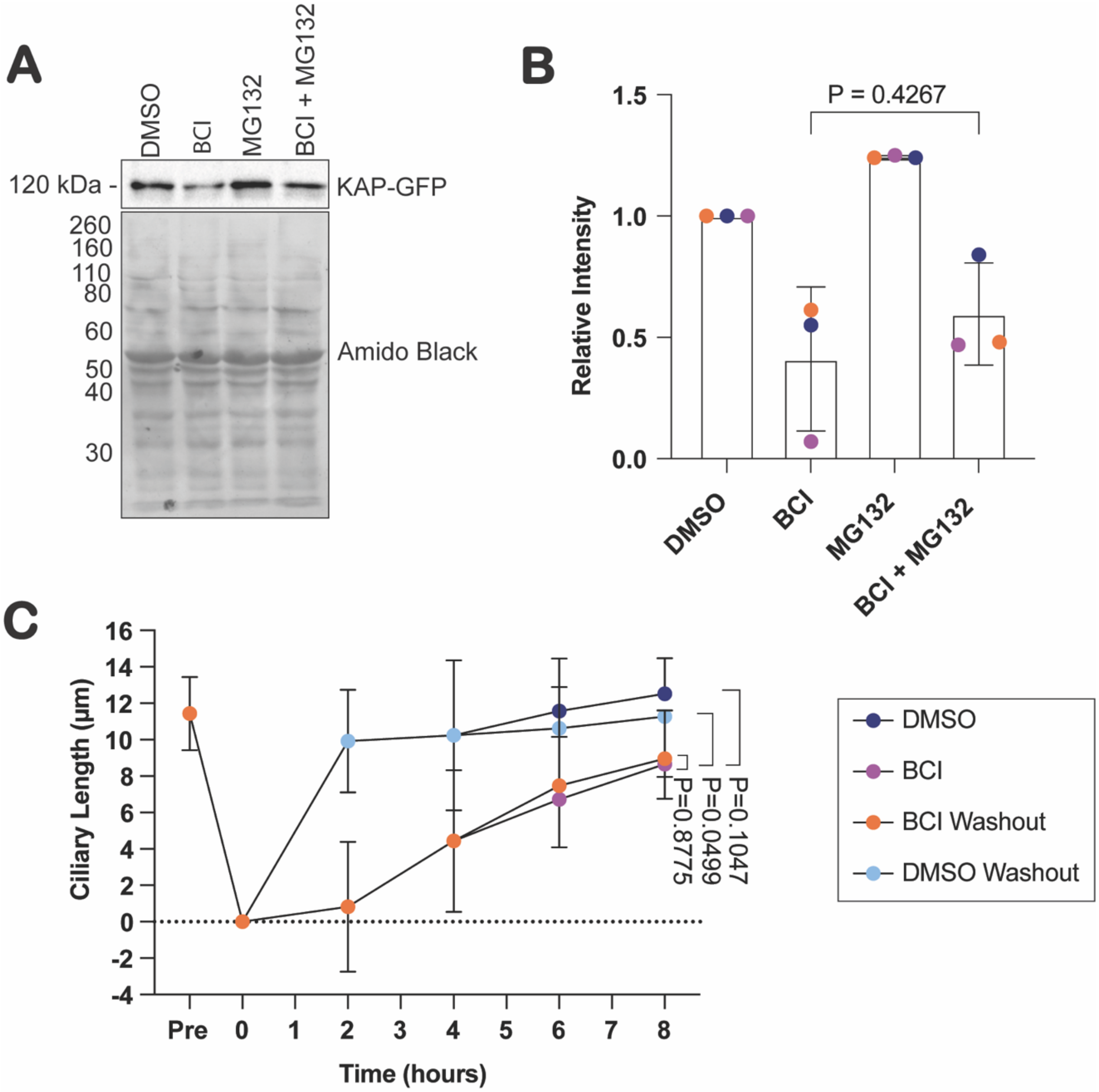


**Supplemental Figure 2. BCI inhibits protein synthesis.** (**A**) Western blot of KAP-GFP expression compared to total protein exposed with amido black. Cells were treated for 2 hours with either 0.5% DMSO, 30 $\mu$M BCI, 50 $\mu$M MG132, or both 50 $\mu$M MG132 and 30 $\mu$MBCI. (**B**) Quantification of (A). Error bars are standard deviation of the mean (n=1, N=3). The P value was determined by an unpaired t-test between BCI and BCI+MG132. **(C)** Single regeneration performed in parallel with Figure 3H. Error bars are mean with 95% confidence interval (n=30, N=3). P values were determined with a two-way ANOVA with Tukey's correction for multiple comparisons.

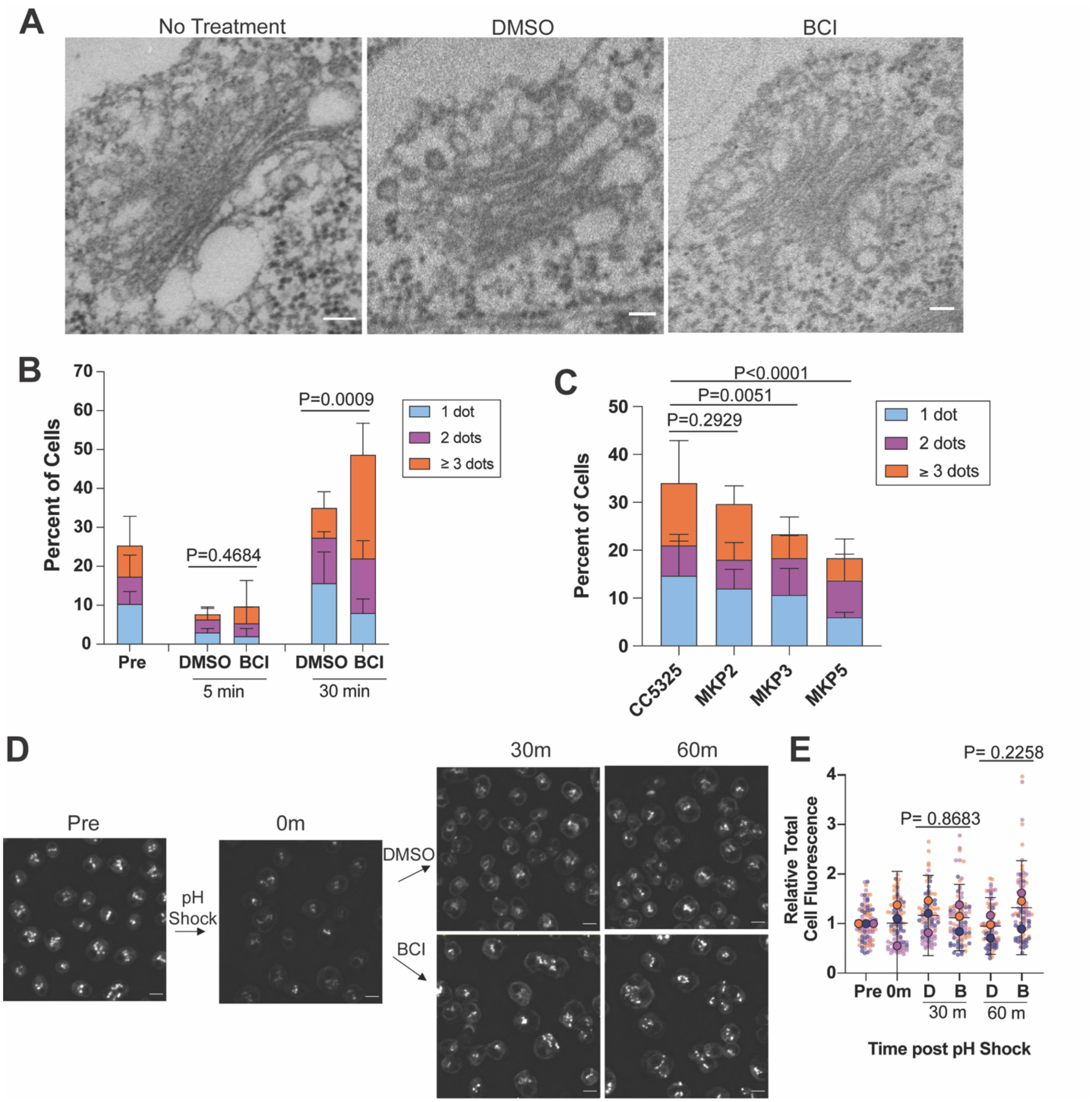


**Supplemental Figure 3.** ***Chlamydomonas*** **DUSP6 orthologs and tagged Arl6 show membrane trafficking defects.** (**A**) Representative EM images of the Golgi apparatus in cells with no treatment, 0.5% DMSO, or 30 $\mu$M BCI. Scale bars are 100nm. (**B**) Quantification of actin puncta in regenerating wild type cells in either BCI or DMSO. Error bars are mean with standard deviation (n=100, N=3). P values were determined using an unpaired t-test. (**C**) Quantification of actin puncta in steady state DUSP6 orthologs. Error bars are mean with standard deviation (n=100, N=3). P values were determined using a Chi-square test (P<0.0001) and a two-sided Fisher's exact test comparing no dots to total dots. Individual bars represent average percent of cells (n=100, N=3). (**D**) Representative images of Arl6/F02 venus-tagged *Chlamydomonas* cells regenerated in 0.5% DMSO or 30 $\mu$M BCI for 1 hour. (**E**) Quantification of arl6 whole cell fluorescence. Error bars are mean with 95% confidence interval (n=30, N=3). P values were determined using an unpaired t test.

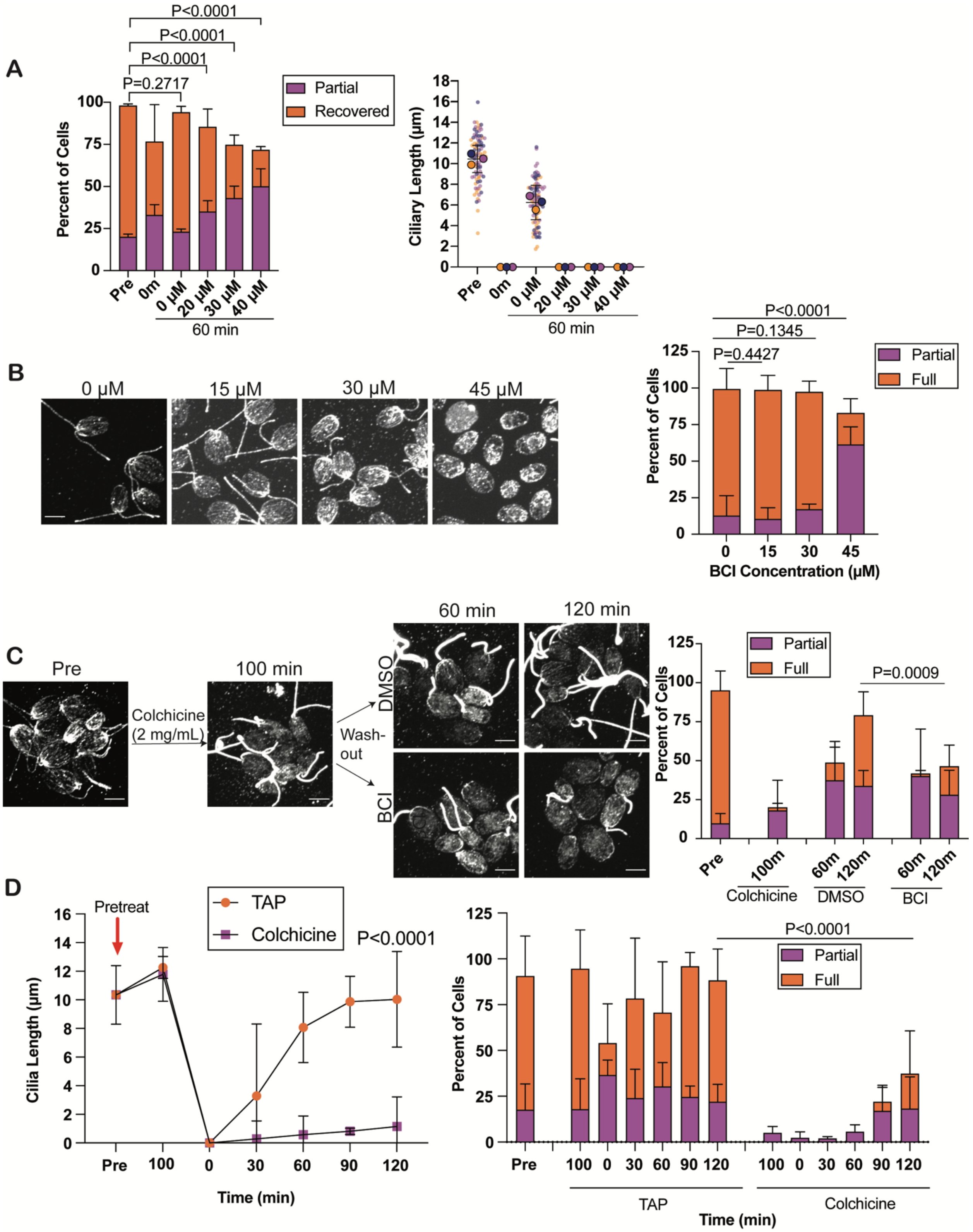


**Supplemental Figure 4. Initial colchicine-induced depolymerization prevents ciliogenesis and microtubule reorganization.** (**A**) Cells were regenerated in either 0.4% DMSO or increasing concentrations of BCI and stained for B-tubulin. Error bars are the mean with standard deviation for the microtubule quantification (left) (n=100, N=3). P values for microtubule reorganizaiton were determined using a Chi-square test with a two-sided Fisher's exact test. The ciliary length quantification (right) shows the mean with the 95% confidence interval (n=30, N=3). (**B**) Steady state wild type cells were treated with either 0.5% DMSO or increasing concentrations of BCI for 2 hours and stained for B-tubulin. Error bars are mean with standard deviation (n=100, N=3). P values were determined with a Chi-square test ($p<0.0001$) and a two-sided Fisher's exact test. (**C**) Cells were pretreated with 2 mg/mL colchicine for 100 minutes. Colchicine was washed out and cells were incubated with either 0.2% DMSO or 20 $\mu$M BCI immediately following. Cells were stained for B-tubulin. Microtubule repolymerization is quantified on the right. Error bars are mean with standard deviation (n=100, N=3). P values were determined using a Chi-square test ($p=0.0009$) and two-sided Fisher's exact test for the 120 minute time points. (**D**) Cells were pretreated with either TAP or 2 mg/mL colchicine for 100 minutes and then regenerated in TAP. Cells were fixed and stained for B-tubulin. For ciliary length measurements, error bars are mean with 95% confidence interval (n=30, N=3). The P value for the 120 minute ciliary timepoints was determined using a two-way ANOVA with Tukey's correction. Microtubule repolymerization is quantified on the right. Error bars are mean with standard deviation (n=100, N=3). P values for the 120 minute microtubule reorganization were determined using a Chi-square test ($p=0.0008$) and two-sided Fisher's exact test.

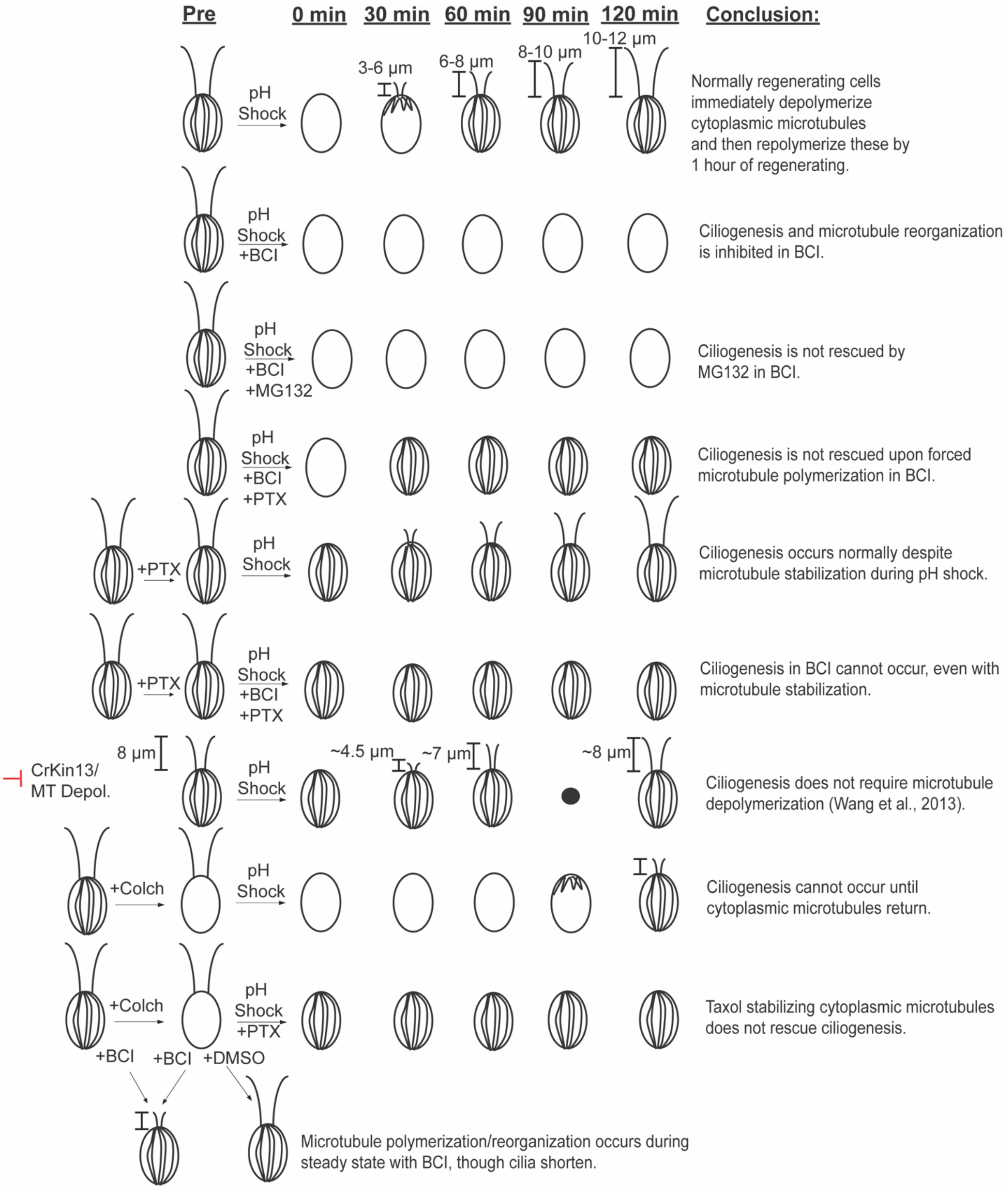


**Supplemental Figure 5. Summary of microtubule data.**

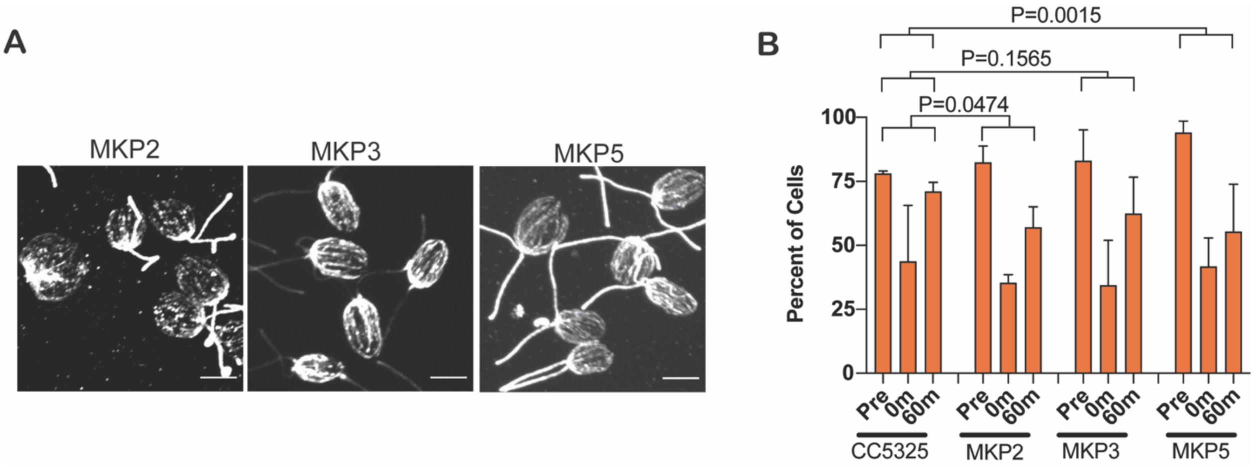


**Supplemental Figure 6.** ***Chlamydomonas*** **DUSP6 orthologs exhibit microtubule polymerization defects.** (**A**) *Chlamydomonas* DUSP6 orthologs were regenerated and stained for B-tubulin. Scale bars are 5 $\mu$m. (**B**) The mean number of cells over 3 trials is plotted with standard deviation (n=100, N=3). P values compare the pre and 60 minute full microtubule quantifications between CC5325 and each mutant. The P values were calculated using a Chi-square test (P=0.0136) with a two-sided Fisher's exact test.
